# Deep learning based design of buried hydrogen bond networks with HBDesigner

**DOI:** 10.64898/2026.06.08.730848

**Authors:** Henry Dieckhaus, Brock T. Harvey, Tomiris Mulikova, Jessica T. Horenstein, Nathan I. Nicely, Nicholas Z. Randolph, Brian Kuhlman

**Affiliations:** Department of Biochemistry and Biophysics, School of Medicine, University of North Carolina at Chapel Hill, Chapel Hill, North Carolina, USA; Division of Chemical Biology and Medicinal Chemistry, Eshelman School of Pharmacy, University of North Carolina at Chapel Hill, Chapel Hill, North Carolina, USA; Department of Pharmacology, School of Medicine, University of North Carolina at Chapel Hill, Chapel Hill, North Carolina, USA; Department of Bioinformatics and Computational Biology, School of Medicine, University of North Carolina at Chapel Hill, Chapel Hill, North Carolina, USA; Lineberger Comprehensive Cancer Center, School of Medicine, University of North Carolina at Chapel Hill, Chapel Hill, North Carolina, USA

**Author notes:** these authors contributed equally to this work (Henry Dieckhaus, Brock T. Harvey, Tomiris Mulikova).

## Abstract

Accurate design of hydrogen-bonding (H-bonding) interactions is a longstanding goal in protein design, as they can facilitate specific protein-protein interactions while improving the solubility of the proteins in the unbound state. Despite this, computational design of H-bond networks remains underexplored in the deep learning era. Here, we present HBDesigner, a novel algorithm for H-bond network design. Through a combination of deep learning-based sampling and atomistic energy scoring, HBDesigner outperforms existing tools in designing connected H-bond networks onto protein scaffolds. We demonstrate the usefulness of HBDesigner by creating monomeric proteins with buried polar interactions and homodimers with extended interface H-bond networks, and by installing specificity into a family of homologous heterodimers where prior design tools fail to do so. The ability to design H-bond networks into arbitrary protein scaffolds should be broadly useful for a wide range of design applications.

## Introduction

Hydrogen bonds (H-bonds) play a critical role in dictating protein structure and function. Enzymes use H-bonds to preorganize key groups for catalysis, enhancing efficiency by reducing the entropic cost of binding^1^. Binding affinity^2^ and specificity^3^ of protein-protein interactions can also be driven by H-bonding contacts, which require precise geometric compatibility between subunits^4^. For minibinders and protein-based nanoparticles, increasing interface polarity can enhance solubility and reduce aggregation^5^.

Despite these factors, tools for de novo protein design have historically struggled to design protein pockets and interfaces with H-bonding networks comparable in complexity to those observed in naturally occurring proteins^4,6^. This is a challenging design problem because energetically favorable H-bonds require accurate geometric placement^7^ and sufficient compensation for desolvation^8^. Even in the era of deep learning (DL) based protein design, most de novo binder design campaigns still rely on hydrophobic contacts to achieve strong experimental success rates^4,9,10^. This strategy can limit the scope of viable targets, while many therapeutically relevant target classes such as antigen-antibody complexes exhibit polar interfaces^11^.

Previous efforts to design buried H-bond networks led to the Rosetta HBNet protocol^12^, which searches for combinations of sidechain rotamers capable of H-bonding when installed onto a given backbone. This approach has been successfully applied to several de novo systems, including pH-sensitive assemblies^13^ and transmembrane β-barrels^14^. A later update replacing exhaustive network enumeration with Monte Carlo sampling (MC HBNet) improved its performance on large, asymmetric systems^8^, facilitating the design of orthogonal heteromeric interfaces^15^. While effective, MC HBNet suffers from a few limitations, including a reliance on fixed rotamer libraries and difficulty modeling long, flexible amino acids. More recently, DL design tools such as ProteinMPNN^16^ and PIPPack^17^ have demonstrated significant advancements in sequence design and sidechain packing, respectively, in comparison to empirical methods such as Rosetta. ProteinMPNN in particular has been used to improve the polarity of designed protein-protein interfaces^5^. However, these and other DL design tools lack the atomistic energy function necessary to explicitly score and optimize H-bond geometry toward fine-grained metrics such as saturation or buried unsatisfied polar atoms (BUHs), so they cannot be readily adapted to design H-bond networks with high connectivity or specific amino acid compositions.

We hypothesized that a DL approach could be used to improve the speed and accuracy of H-bond network design. Here, we introduce HBDesigner, a fixed-backbone protein design algorithm trained to install H-bond networks onto protein scaffolds. Our method integrates recent advancements in DL based sequence design and sidechain packing with the atomistic scoring capabilities of the Rosetta energy function. We demonstrate its efficacy at installing designable buried polar networks through several *in silico* benchmarks on diverse sets of native and de novo designed proteins. We then apply our method to design de novo proteins with buried H-bond networks integrated into their cores and interfaces, demonstrating high experimental success rates. Finally, we demonstrate the capability of HBDesigner to enhance binder specificity by designing and validating a family of homologous heterodimers with interface networks that impart specificity into otherwise promiscuous scaffolds.

## Results

### Modeling hydrogen bond networks in silico

We sought an algorithm that receives a protein backbone as input and rapidly samples energetically favorable H-bond networks. To achieve this, we formulated HBDesigner as a three-step procedure **(Fig. 1A).** The first two steps use graph neural networks to encode the backbone structure into a protein graph. The design model **(Fig. 1A, top)** autoregressively predicts both the position and the amino acid identities of only the network residues. This sparse, network-based decoding scheme, which enables rapid sampling, is distinct from inverse folding models such as ProteinMPNN, which decode only amino acid identities for a pre-specified designable region. The packing model **(Fig. 1A, middle)** then predicts the sidechain chi angles for residues in the network. The third step, the scoring module **(Fig. 1A, bottom)**, uses PyRosetta to rapidly minimize, score, and rank the predicted networks.

**Figure 1:**
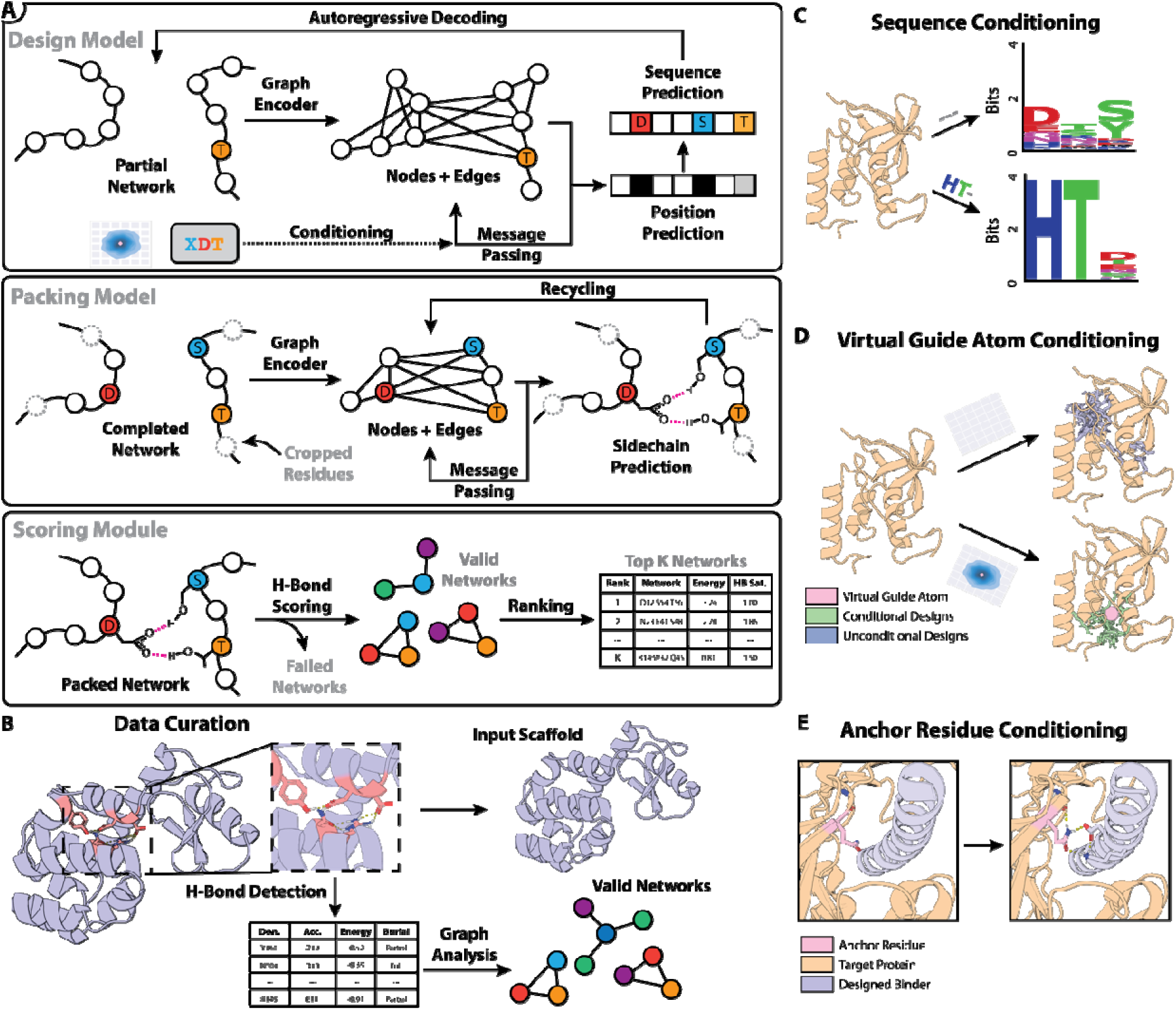
Modeling hydrogen-bonding networks with HBDesigner. **A)** The HBDesigner algorithm consists of three steps. First, the design model encodes the backbone structure into a graph and assigns amino acid identities to a small cluster of residues (top). Second, the packing model predicts sidechain chi angles based on local backbone geometry (middle). Third, the scoring module (bottom) calculates atomistic energy terms and ranks the resulting networks. **B)** H-bonds were identified in native proteins using Rosetta and collected into graphs to represent individual H-bond networks. Pairs of scaffolds and networks were then randomly sampled during training. **C)** Sequence conditioning may be provided to specify desired amino acid composition in the predicted networks. Unconditional sampling (top) produces diverse networks potentially including all polar amino acids, while conditional sampling (bottom) produces networks matching the specified sequence composition. **D)** A virtual guide atom (pink sphere) may be provided to denote the approximate centroid of the desired networks. This can be used to shift the sampling toward a certain region (bottom) which may be neglected by unconditional sampling (top). **E)** One or more fixed anchor residues (pink) may be provided as initial seeds for residue decoding. This enables one-sided design of binder interfaces or extension of existing H-bond networks.

HBDesigner was trained on a dataset of over 1.7 million high-quality native sidechain H-bond networks curated from an August 2021 snapshot of the Protein Data Bank^18^. PyRosetta^19^ was used to optimize and detect networks of H-bonds with favorable geometry to use for training **(Fig. 1B)**. To provide fine-grained control over network design, HBDesigner was trained to use different types of conditioning information, which can be provided at inference time. Sequence conditioning **(Fig. 1C)** specifies the desired amino acid composition of designed networks, while virtual guide atom conditioning **(Fig. 1D)** specifies an approximate center-of-mass for the designed networks in 3D space. The model can also condition on one or more existing polar residues to be used to anchor predicted networks **(Fig. 1E).**

### HBDesigner produces native-like, well-packed networks

We first evaluated HBDesigner by generating H-bond networks for our test set of native protein backbones. When sampled unconditionally, the design model closely reproduced the amino acid distribution of native H-bond networks **(Fig. 2A, blue and gold)**. Generating designs with a low sampling temperature **(Fig. 2A, green)** resulted in a modest increase in small hydroxyl and acidic residues at the expense of aromatic and amidic residues. The generated networks were also native-like in terms of residue co-occurrence, burial, and fraction of sidechain-sidechain vs sidechain-backbone H-bonds **(Extended Data Fig. 1)**. Some higher-order network properties such as saturation, BUHs/BUPHs, and HBNet Score were slightly worse in the designed networks, but this could be mostly mitigated by using a lower sampling temperature **(Extended Data Fig. 1B)**. We next examined the effect of different types of conditioning on HBDesigner predictions. As expected, conditioning on the native network sequence increased native amino acid recovery from 34% to 77%, while providing the approximate native network centroid as a virtual guide atom improved recovery of native network positions from 14% to 43% **(Extended Data Fig. 2A-B)**. Finally, anchor residue conditioning with selected residues from the native network progressively increased recovery of the remaining native residues **(Extended Data Fig. 2C)**. These results indicate the model is responsive to conditioning toward desired network properties.

**Figure 2:**
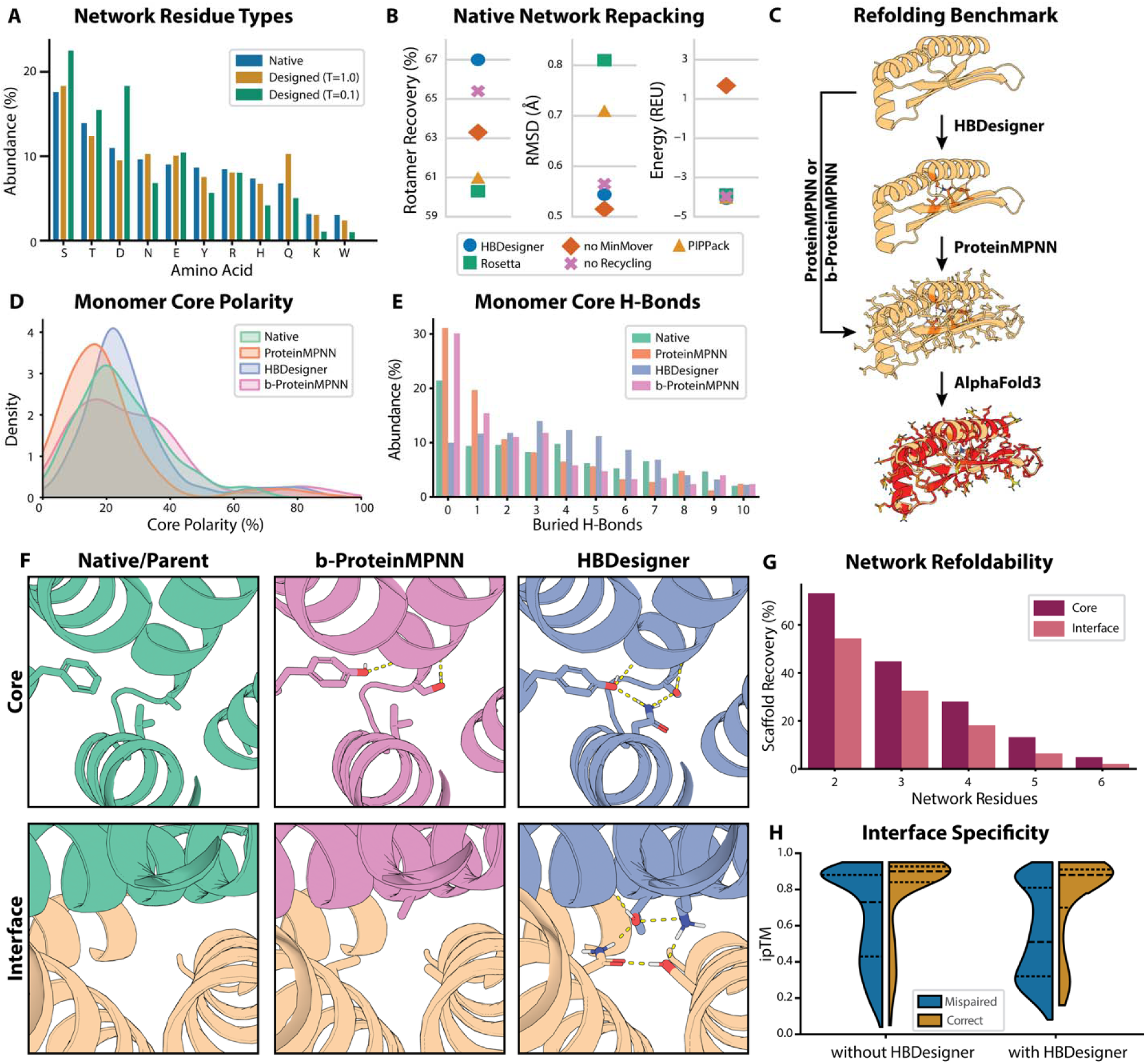
HBDesigner produces native-like, designable hydrogen bond networks when evaluated on a diverse set of unseen scaffolds. **A)** Relative abundance of each polar amino acid in test set native H-bond networks compared to networks generated by the HBDesigner sequence design model with either high (T=1.0) or low (T=0.1) sampling temperature (n=525). **B)** Sidechain packing statistics for different packing models, evaluated by repacking test set native H-bond networks without native sequence context (n=1,011). **C)** Workflow of the refolding benchmark used to evaluate the performance of HBDesigner and other methods. **D)** Percentage of polar core residues found in a diverse set of native protein monomers (n=642), compared to sequences designed for the same backbones using either ProteinMPNN, HBDesigner or b-ProteinMPNN. **E)** Comparison of buried hydrogen bonds involving sidechains detected in native monomers (n=642) and sequences designed by either HBDesigner, ProteinMPNN or ProteinMPNN biased toward buried polar residues (b-ProteinMPNN). **F)** Example of a nonpolar buried pocket in a native monomer (top) and a de novo heterodimer interface (bottom) which were redesigned to be more polar using b-ProteinMPNN (middle) or HBDesigner (right). **G)** Scaffold recovery percentages for designed core and interface H-bond networks of different sizes obtained after sequence design and refolding, evaluated on de novo generated monomer (core, red, n=726) and heterodimer (interface, pink, n=690) backbones. **H)** Distribution of AlphaFold3 interface predicted template modeling (ipTM) scores obtained for de novo heterodimers predicted with subunits correctly paired (gold, n=190) or mispaired (blue, n=1,710) after sequence design with- or without HBDesigner. Dashed lines indicate quartiles for each distribution.

Packing methods were evaluated by repacking native networks onto their original backbones without any surrounding sequence context. The HBDesigner packing model produced higher rotamer recovery and lower root-mean-square deviation (RMSD) than the other two methods tested, the DL-based packing model PIPPack and the Rosetta PackRotamersMover **(Fig. 2B, left and middle)**. Rotamer recovery and RMSD were slightly improved by the use of recycling **(Fig. 2B, left and middle, pink)**, a procedure in which predicted sidechains are “recycled” back through the model multiple times to allow further refinement. Meanwhile, postprocessing with the Rosetta MinMover was essential to avoid clashing and disfavored Rosetta energy scores **(Fig. 2B, right, orange)**. This also improved rotamer recovery at the cost of a slight increase in RMSD.

### HBDesigner effectively improves designed protein polarity in silico

To evaluate HBDesigner in the context of an actual design campaign, we ran a refolding benchmark on a large library of native and de novo generated backbones **(Fig. 2C)**. In brief, we designed sequences for each backbone using either a) HBDesigner followed by ProteinMPNN to design the remaining sequence, b) ProteinMPNN alone, or c) ProteinMPNN with amino acid biasing enabled to encourage buried polar residues (referred to as b-ProteinMPNN). We then refolded all sequences using AlphaFold3^20^ (AF3), then minimized and scored all successfully refolded designs with PyRosetta to assess their polarity and H-bonding. On a test set of 532 native monomer backbones, we found that sequences designed with ProteinMPNN had significantly less polar cores after refolding than those of the native proteins **(Fig. 2D, green and orange).** Generating even small (3-residue) buried networks with HBDesigner prior to design with ProteinMPNN resulted in a marked increase in core polarity **(Fig. 2D, purple)** and an accompanying increase in buried H-bonds **(Fig. 2E, purple and orange)**. Importantly, simply increasing the core polarity using amino acid biasing generated fewer buried H-bonds than HBDesigner **(Fig. 2E, pink)**, indicating that our method was indeed yielding designs enriched with productive H-bonding motifs. Upon visual inspection, we observed that HBDesigner more often generated highly connected networks with branched interactions, rather than the isolated H-bonds common to b-ProteinMPNN **(Fig. 2F, top)**. In other cases, HBDesigner installed a novel network into a pocket that b-ProteinMPNN would otherwise keep nonpolar **(Fig. 2F, bottom)**. These findings were replicated on an additional set of 726 de novo monomer backbones **(Extended Data Fig. 3A)**, as well as 690 de novo heterodimers **(Extended Data Fig. 3B)**, where HBDesigner produced more buried core or interface H-bonds, respectively, than the other methods.

Using the same refolding test, we also compared HBDesigner to the Rosetta MC HBNet protocol. We applied each method to design 5 buried networks per scaffold for our libraries of de novo monomer and heterodimer backbones and evaluated them with various metrics, including *saturation*, which is the fraction of total H-bond donor/acceptor sites in a given set of residues that are engaged in H-bonding^8^. We then designed sequences around each network with ProteinMPNN and assessed their refoldability by AF3 pLDDT. For each method, we also calculated *scaffold recovery percentage*, which is the percentage of backbones in the library with a designed H-bond network that was fully recapitulated after refolding. We found that the two methods produce similar results when used to design monomer cores **(Table 1, top)**, with HBDesigner producing better AF3 pLDDT and saturation but MC HBNet achieving better scaffold recovery and finding more networks. When used to design heterodimer interfaces **(Table 1, bottom)**, however, HBDesigner achieves slightly better scores on all metrics.

**Table 1:** Performance comparison of HBDesigner and MC HBNet. Runtimes are reported as mean ± standard deviation for each method. The best score reported on each dataset is bolded for each metric.

| De Novo Monomers |  |  |  |  |  |
| --- | --- | --- | --- | --- | --- |
| Method | Runtime (s) | Networks Found | Top-5 Saturation | AF3 pLDDT | Scaffold Recovery (%) |
| HBDesigner | 7.6 ± 0.6 | 8.9 | **0.69** | **79** | 45 |
| MC HBNet | **5.5 ± 1.9** | **13.5** | 0.65 | 76 | **51** |
| De Novo Heterodimers |  |  |  |  |  |
| Method | Runtime (s) | Networks Found | Top-5 Saturation | AF3 ipTM | Scaffold Recovery (%) |
| HBDesigner | **9.3 ± 0.7** | **11.2** | **0.72** | **0.59** | **32** |
| MC HBNet | 28 ± 6 | 7.7 | 0.69 | 0.58 | 29 |
| MC HBNet* | 7 ± 2 | 5.2 | 0.68 | 0.58 | 22 |

\* Run with a more restrictive MoveMap only allowing interface residues to be designed.

We next sought to characterize the upper limits of H-bond network design with HBDesigner and AF3. To do this, we repeated the above refolding test and generated designs including buried networks of 2-6 residues, then assessed their scaffold recovery scores as a function of network size. We observed that networks designed across heterodimer interfaces were less recoverable than those designed into monomer cores **(Fig. 2G, red and pink bars)**. We also found that scaffold recovery decreased sharply with increasing network size, as over 70% of monomer scaffolds could accommodate a 2-residue network in the core, compared to less than 4% for 6-residue networks **(Fig. 2G, red bars)**.

Another potential use for HBDesigner is to design second-shell interactions around functional sites. To assess this capability, we ran HBDesigner on several dozen metal-binding sites curated from MetalPDB^21^, along with a similar number of native enzyme active sites from the AME benchmark dataset^22^, with a subset of metal-/ligand-contacting residues provided to the model as anchor residues **(Extended Data Fig. 4)**. We observed that, among its top-10 ranked designs, HBDesigner recovered 29% of second-shell interactions in the MetalPDB test set, and a slightly higher 39% of native residues in the AME dataset. These results suggest that our model can be applied to design second-shell interactions to stabilize existing functional sites.

### HBDesigner improves predicted binding specificity in otherwise promiscuous scaffolds

Next, we wished to test our hypothesis that installing H-bond networks into protein-protein interfaces with HBDesigner could impart improved binding specificity. We collected 19 de novo designed heterodimers from prior design efforts and generated a family of 10 highly similar backbones (template modeling [TM] score>0.9) for each scaffold using partial RFdiffusion^23^. We designed sequences for each backbone using ProteinMPNN, with and without first installing a 4-residue network across the interface using HBDesigner. We then generated all possible A/B subunit pairs within each family and refolded them with AF3, using the interface predicted template modeling (ipTM) confidence score as a proxy for predicted binding. When designed only with ProteinMPNN, the correctly paired dimers received high ipTM scores (median=0.90), but the mispaired dimers also received moderate ipTMs (median=0.73), indicating only modest specificity for the designed dimer interactions **(Fig. 2H, left)**. When using HBDesigner, the correctly paired dimers produced only slightly lower ipTMs (median=0.88), while the mispaired dimers received significantly worse scores (median=0.51), indicating higher predicted specificity **(Fig. 2H, right)**. When stratified by family, HBDesigner interfaces show higher predicted specificity (ΔipTM between correct and mispaired dimers) than ProteinMPNN for 13/19 families **(Extended Data Fig. 5A)**. We also plotted ΔipTM against structural characteristics of each family, and we observe a general trend toward lower ΔipTMs for larger complexes with greater similarity and more interface contacts **(Extended Data Fig. 5B-D)**. However, due to the small sample size of this analysis (n=19), these effects have wide confidence intervals and low statistical power.

### Design and characterization of de novo monomers with polar cores

As an initial experimental test of our model, we designed a set of small (80-120 residue), de novo monomeric proteins, using HBDesigner to install a single H-bond network of 3-6 residues into the core of each protein. We selected 19 designs with a range of buried polar surface area percentages (bpSA%) for expression and characterization by circular dichroism (CD). Of these, 11 were found to be well-expressed, stable, and folded with a CD curve consistent with the designed fold **(Fig. 3A-C, Extended Data Fig. 6)**. Notably, the successful designs included a diverse range of network topologies and amino acid compositions. Helical bundles had the highest expression success rate (9 of 11 designs), while mixed α/β folds were moderately successful (2 of 4), and all β-sheet-rich designs (0 of 4) failed **(Fig. 3D)**. While designs with the highest bpSA% were the least likely to express well, they also typically had the most β-sheet character (**Fig. 3D)**. We also observed that seven of the twelve purified designs were folded up to >95°C **(Fig. 3C, Extended Data Fig. 6)**, despite having significantly more polar cores than typical designs produced by ProteinMPNN **(Fig. 3D)**.

**Figure 3:**
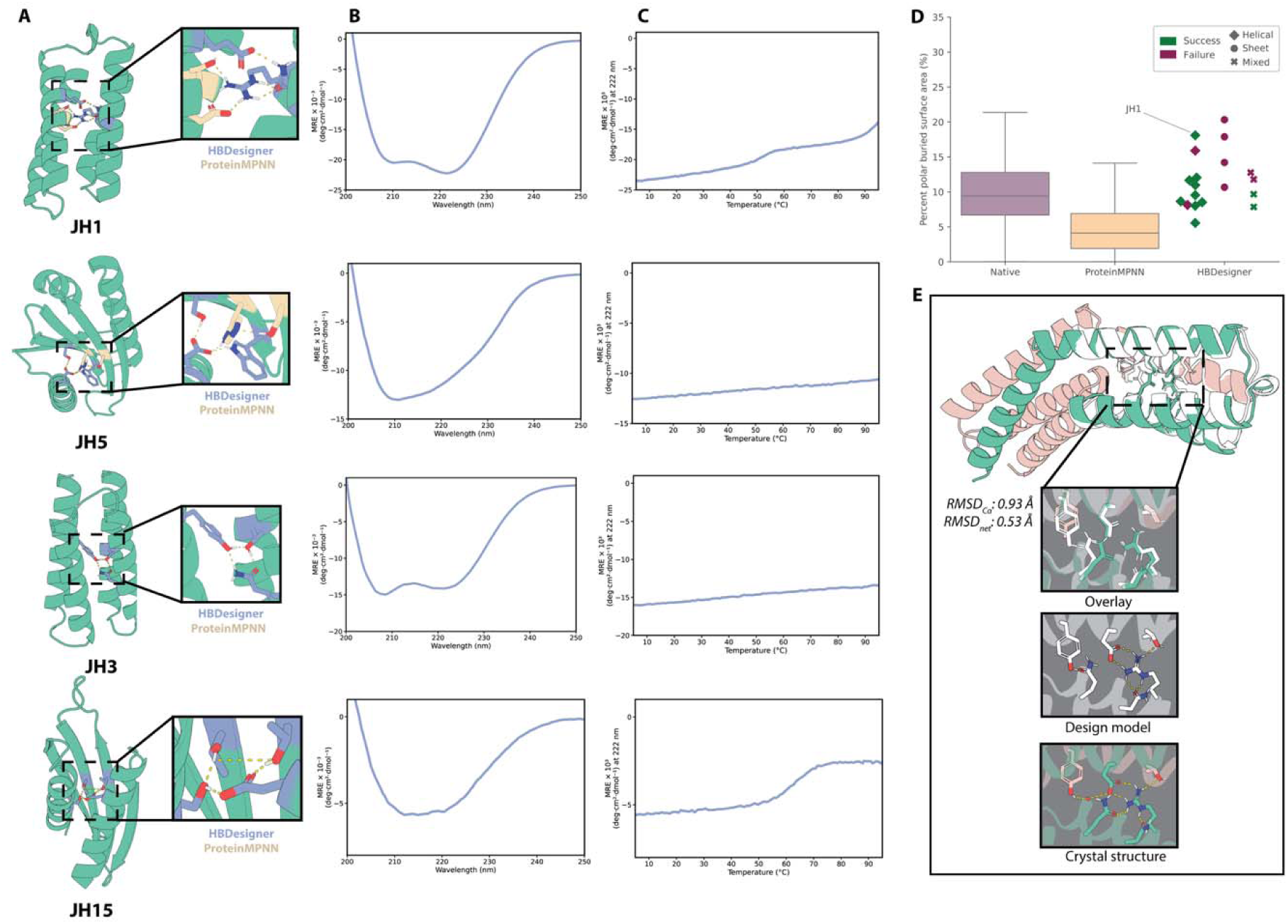
De novo design and validation of monomers with highly polar cores. **A)** Design models of selected successful monomer designs with different folds and network geometries. Insets show designed H-bond networks comprised of residues introduced by HBDesigner (blue) and ProteinMPNN (tan). **B)** Circular dichroism (CD) spectra and **C)** temperature melts of the selected monomers. **D)** Core polarity, calculated as the percentage of buried surface area originating from polar atoms, for the tested designs (n=19), native monomers (n=642), and ProteinMPNN designs (n=629). Designs that could be expressed, purified, and validated by CD were considered successful. Marker shapes denote the fold type of each design as assessed by visual inspection. **E)** Comparison of the design model (white) and crystal structure (green) for design **JH1** (PDB: 13JM, 2.1 Å). The second chain of the crystal structure is colored in taupe. Structural waters are shown as red spheres and the aligned root-mean-squared deviation of the Cα atoms (RMSD_Cα_) and the network sidechain heavy atoms (RMSD_net_) are reported.

We solved the crystal structure for Design **JH1** (PDB: 13JM, 2.1 Å resolution), a four-helix bundle with an extended network centered around a buried arginine **(Fig. 3A, top)**. Under crystallization conditions, the protein forms a domain swap dimer in which the loop connecting helix 1 and 2 forms an extended conformation that results in helix 1 from chain A partnering with helices 2-4 of chain B **(Fig. 3E)**. In solution at a lower concentration, **JH1** exists in a monomer-dimer equilibrium as evidenced by size exclusion chromatography **(Extended Data Fig. 7A)**. Aside from the alternate conformation observed for loop 1, each four-helix unit in the crystal structure closely aligns with the design model (RMSD_Cα_ = 0.93 Å, RMSD_net_ = 0.53 Å), allowing a direct comparison of the designed and experimentally determined H-bond networks **(Fig. 3E)**. As designed, R86 is buried in the protein core and forms H-bonds with the sidechains of E36, N63, and Q66, as well as T10 coming from the helix on the opposite chain. One interaction in the crystal structure that is not captured in the design model is a structural water that bridges between Y17 on helix 1 and the designed hydrogen bond network **(Fig. 3E, bottom)**. Interestingly, AF3 predicts that the protein will domain swap when predicted as a dimer, potentially reflecting strain in loop 1 when the protein folds into a monomer **(Extended Data Fig. 7B).** Based on this hypothesis, we redesigned loop 1 to reduce strain in the monomer state and mitigate domain swapping. From this effort, we obtained a new set of designs which AF3 no longer predicted to dimerize or domain swap **(Extended Data Fig. 7C-D)**, with a reduced dimer ipTM of 0.25 compared to 0.89 for JH1. Notably, the polar core of a protein remained intact, suggesting that additional design and filtering steps can reduce dimerization propensity while maintaining the originally designed core.

For our remaining designs which expressed but failed to crystallize (n=10), we ran *in silico* tests to further validate our predicted structures **(Extended Data Fig. 8A)**. First, we refolded each design with three structure predictors not used for filtering and observed that nearly all designs adopted highly similar folds (median TM score > 0.95, median RMSD_Cα_ < 0.8Å for each method), with the predicted network sidechains closely recapitulating the design model (median RMSD_network_ < 0.9 Å). Second, we used PLACER^24^, a fixed-backbone denoising model, to predict the sidechains of each network and its nearest neighbors packed onto the original design model backbone. Again, we observed close agreement with the designed network sidechains (median RMSD = 0.26 Å) and high confidence (median pLDDT = 0.97) in all cases. These results across several orthogonal methods further support our hypothesis that our designs are likely to adopt the intended fold and network geometry.

### Design and characterization of de novo homodimers with highly polar interfaces

To examine the ability of HBDesigner to design interfaces with buried hydrogen bonds, we generated de novo homodimer backbones of 200-400 amino acids and used HBDesigner to introduce H-bond networks consisting of two, three, or four polar residues across each interface. Rather than installing a single symmetric network, we instead chose to design asymmetric networks and symmetrize them across the interface, meaning that two copies of each network are present in each interface **(Fig. 4A)**. This enabled us to more easily cover a large portion of the interface with our networks. We expressed 22 homodimers and characterized them with mass photometry (MP) to determine their propensity to form dimeric complexes **(Fig. 4C, Extended Data Fig. 9)**. The proteins were produced as fusions with maltose-binding protein to allow easy detection with MP. All 22 designs expressed well, and 15 were found to be primarily dimeric by mass photometry at a concentration of 50 nM. Among the remaining designs, three were a mixture of monomer and dimer, and one showed additional peaks at higher mass **(Extended Data Fig. 9)**. Assessment of interface polarity revealed that our designed interfaces were more polar on average than those produced by ProteinMPNN, while still less polar than native homodimers **(Fig. 4B)**. The one exception was design S2H, which had a 62% polar interface by surface area due to additional backbone-backbone H-bonds from extensive β-strand pairing **(Extended Data Fig. 9)**.

**Figure 4:**
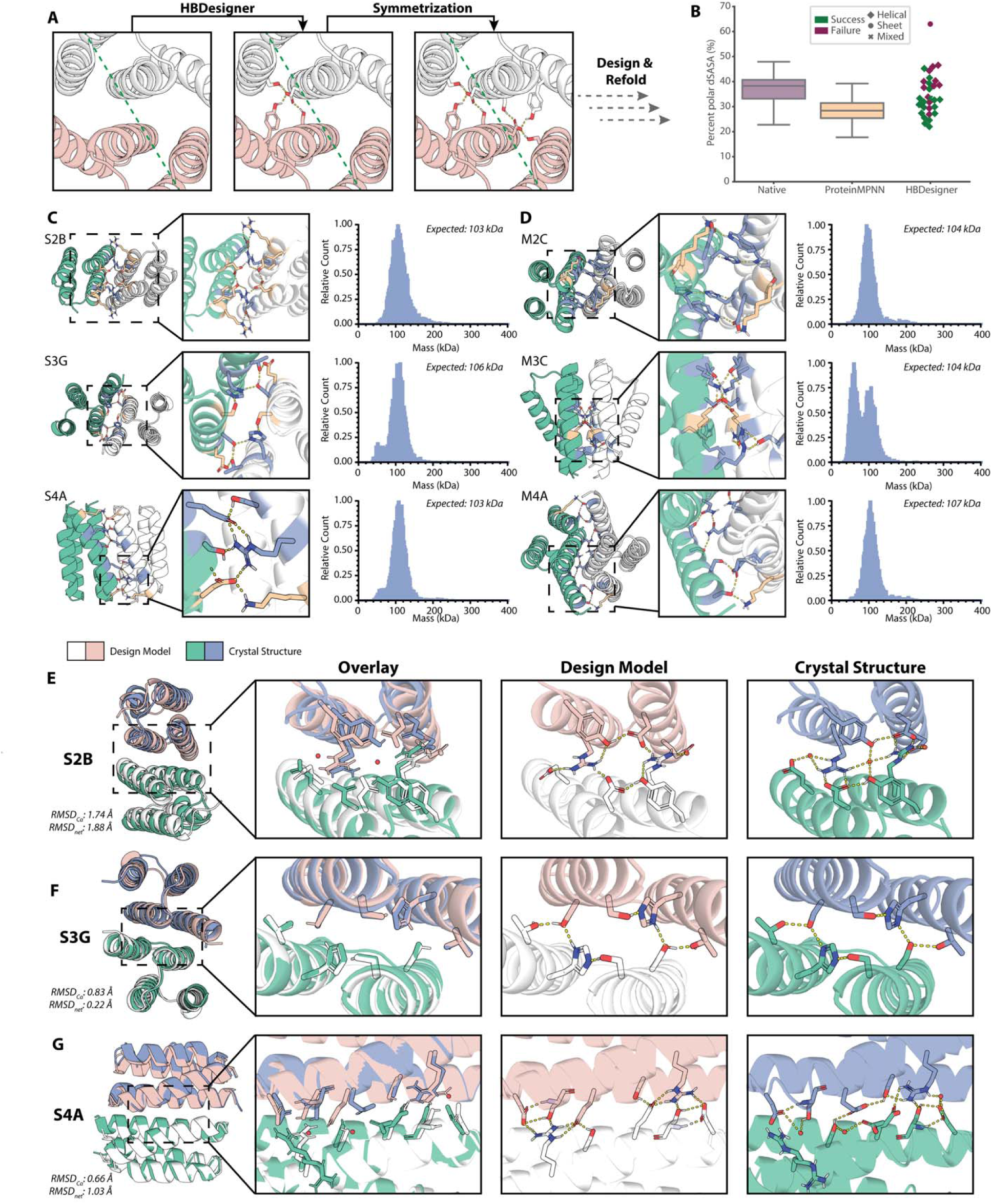
De novo designed obligate homodimers with highly polar interfaces. **A)** Workflow for two-sided homodimer interface design using HBDesigner. Networks were generated by HBDesigner, then symmetrized by copying the network across the asymmetric unit prior to downstream design and refolding. **B)** Interface polarity, calculated as a percentage of total change in solvent-accessible surface area upon binding, for the tested designs (n=31), native obligate homodimers (n=47), and ProteinMPNN designs (n=92). Designs were considered successful if measured as >50% dimeric by mass photometry. Marker shapes denote the fold type of each design as assessed by visual inspection. **C-D)** Design models (left) and mass photometry (MP) plots (right) for selected homodimer designs with **C)** single and **D)** multiple H-bond networks. Insets (middle) show designed H-bond networks comprised of residues introduced by HBDesigner (blue) and ProteinMPNN (tan). MP plots are labeled with expected mass for each protein expressed as an MBP fusion. **E-G)** Comparison of design models with crystal structures obtained for homodimer designs S2B (PDB: 35SL, 1.80 Å), S3G (13FC, 1.79 Å), and S4A (35SB, 1.83 Å). The design models are shown in white and pink, and the crystal structures are shown in green and blue. Structural water molecules are shown as red spheres. The aligned root-mean-squared deviation of the Cα atoms (RMSD_Cα_) and the network sidechain heavy atoms (RMSD_net_) are reported for each design.

Encouraged by the high success rate of our initial campaign in terms of generating stable dimers, we next sought to design even more polar homodimer interfaces. To do this, we combined successfully predicted single-network designs with alternate HBDesigner networks for each corresponding backbone. We selected nine designs from the resulting multi-network candidates. Again, all of the designs expressed well. When analyzed by mass photometry **(Fig. 4D, Extended Data Fig. 10)**, we observed that three of our designs were mostly dimeric, three were mostly monomeric, and three were in equilibrium between monomer and dimer. Across the single- and multi-network campaigns, we observed that more polar interfaces were less likely to be successfully expressed and assembled as stable dimers **(Fig. 4B)**.

We solved high-resolution crystal structures for three of our homodimer designs, S2B (PDB: 35SL, 1.80 Å resolution), S3G (13FC, 1.79 Å), and S4A (35SB, 1.83 Å) **(Fig. 4E-G)**. All three structures show close agreement with the design models in terms of overall fold and interface orientation. In design S2B **(Fig. 4E)**, the network topology remained mostly intact, with minor rearrangements in sidechain geometry and H-bond donor/acceptor pairing (RMSD_Cα_ = 1.74 Å, RMSD_net_ = 1.88 Å). These changes were enabled by a small shift of the C-terminal helix of each subunit toward the opposite chain, as well as two structural waters integrated into the designed network. One of these was found bridging the central Y43 and R47 residues, and one between R47 and solvent-exposed E72. The crystal structure of design S3G **(Fig. 4F)** showed nearly perfect agreement with the design model (RMSD_Cα_ = 0.83 Å, RMSD_net_ = 0.22 Å), including correct recovery of all network residue rotamers. This network, which was more linear and comprised of less flexible residues than that of S2B, showed no structural water intrusions. Design S4A **(Fig. 4G)** included a five-residue network with partial solvent exposure and several longer, more flexible sidechains. In the crystal structure, there are four homodimers per asymmetric unit, and alternative conformations are observed for the network in the four homodimers. The network formed between chain D and chain C most closely resembles the design model, with R37 chain D burying its guanidinium group and forming hydrogen bonds with S27 chain C and E30 chain C, as designed (RMSD_Cα_ = 0.66 Å, RMSD_net_ = 1.03 Å).

As we did for our monomer designs, we ran our remaining well-behaved homodimers (n=15) through additional structure prediction and PLACER analysis **(Extended Data Fig. 8B)**. We again observed high consistency between the predicted structures and our original design models (median TM score > 0.97, RMSD_Cα_ < 0.8 Å, RMSD_network_ < 1.3 Å for each method). PLACER sidechain analysis again showed high predicted alignment with the designed network and high confidence (median RMSD = 0.34 Å, median pLDDT = 0.98). Together, these results suggest our designs are likely adopting the intended fold and interface orientation.

### Design and characterization of de novo heterodimers with improved binding selectivity

Finally, encouraged by the results of our *in silico* specificity test **(Fig. 2H)**, we designed an experiment to assess whether installing H-bond networks across a designed interface could impart binding specificity among otherwise promiscuous binding partners. To do this, we reanalyzed the heterodimers from the *in silico* test to identify families of heterodimers with high predicted specificity when designed with HBDesigner followed by ProteinMPNN **(Fig. 5B, top)** and low predicted specificity when designed with ProteinMPNN alone **(Fig. 5B, bottom)**. From this, we identified a set of 6 homologous heterodimers **(Fig. 5A)**, which we expressed and tested for binding using ELISA. As predicted from AF3 crossdocking, the ProteinMPNN heterodimers showed high binding affinity for both their intended design partners and their off-target partners **(Fig. 5C, bottom)**. The HBDesigner heterodimers, on the other hand, had significantly improved specificity for their intended design partners, at the cost of diminished on-target affinity **(Fig. 5C, top)**. Our best-performing design, HBDes3, showed 110 nM on-target affinity with 100-fold specificity over its cognate off-target partners. These results indicate that HBDesigner and AF3 can be used to explicitly design specific binding between protein subunits with highly similar folds and interface shapes.

**Figure 5:**
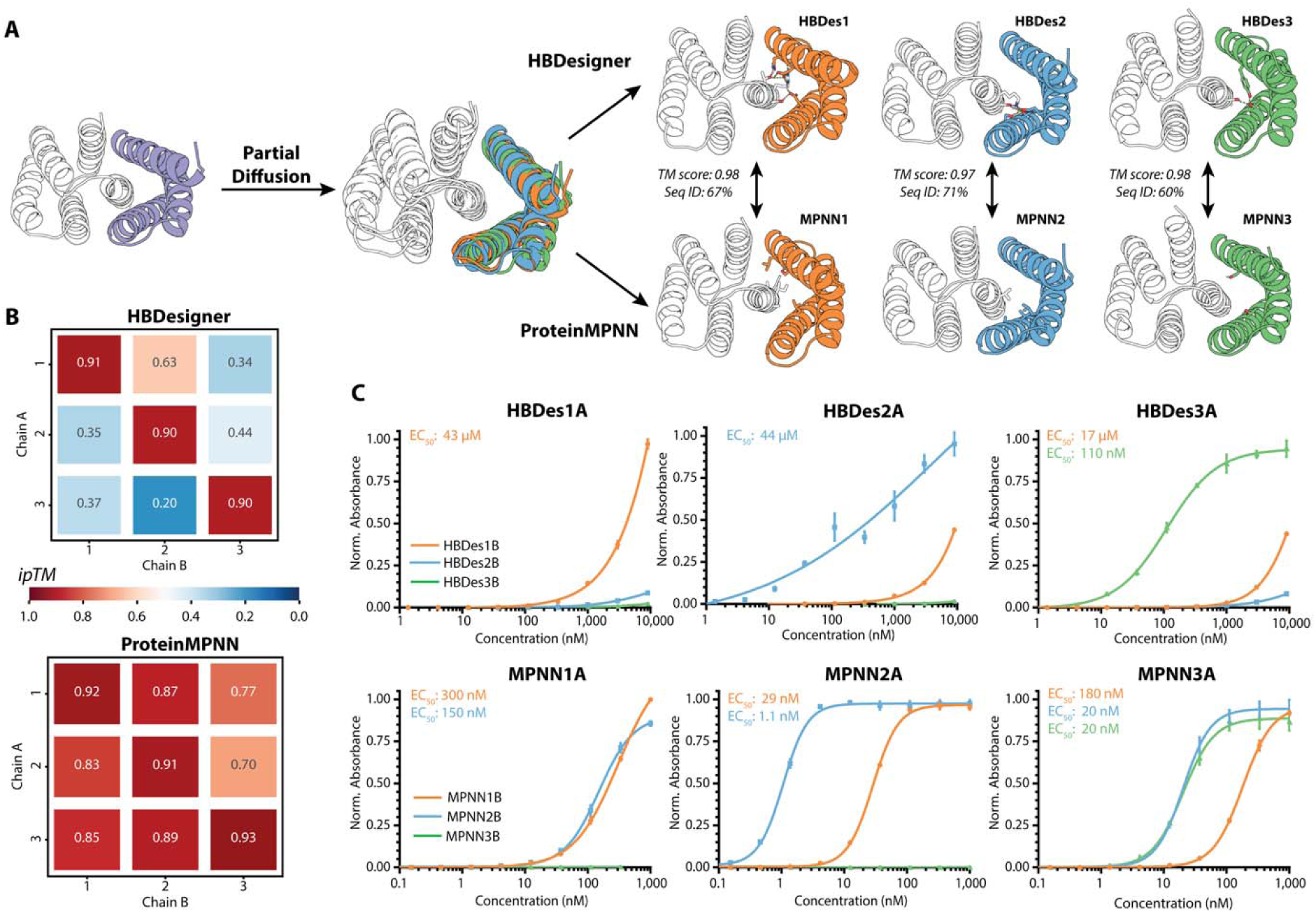
De novo design and validation of heterodimeric proteins with enhanced subunit pairing specificity. **A)** Workflow used to generate structurally similar heterodimers using either HBDesigner (top) or ProteinMPNN (bottom). Design models shown are AlphaFold3 predictions with hydrogens optimized by Rosetta. **B)** Heatmaps showing predicted interface confidence (ipTM) for each combination of heterodimer subunits for the HBDesigner (top) and ProteinMPNN (bottom) sets of designs. **C)** Binding curves measured by ELISA (n=3) for the HBDesigner (top) and ProteinMPNN (bottom) sets of designs. Each subplot is labeled with the identity of the immobilized “A” subunit, while the curve colors indicate the identity of the “B” subunit.

## Discussion

This study was motivated by the hypothesis that a task-specific deep learning (DL) approach to hydrogen-bonding network design could offer advantages over both empirical methods and generalist DL methods by combining aspects of each. In this work, we have demonstrated that this approach, which we call HBDesigner, enables rapid design of H-bond networks into arbitrary protein backbones. On an *in silico* benchmark of over 2000 native and de novo backbones, our method produced buried H-bond networks more reliably than ProteinMPNN while offering similar or better speed and designability compared to MC HBNet. Our experimental results further validate our method by showing how HBDesigner can be readily applied to common design tasks, including installation of H-bond networks into protein cores and both symmetric and asymmetric interfaces.

We developed HBDesigner as a drop-in module intended for use in the early steps of the *in silico* design pipeline. In the experiments described above, we treat H-bond networks as privileged motifs akin to enzyme active sites and follow installation of the network with sequence design for the rest of the protein. In this setting, we observed that ProteinMPNN was often able to extend networks introduced by HBDesigner **(Fig. 3A and Fig. 4C-D, tan sticks)** with additional polar residues. This behavior enabled us to design large (6+ residue) networks starting from seed networks of only 3-4 residues. We also observed that AF3 is capable of modeling H-bonding interactions, with some caveats. While 3 of our 4 crystal structures showed sub-Ångstrom agreement with the design model by Cα RMSD, we often observed subtle sidechain rearrangements, especially at more solvent-exposed positions **(Fig. 3E and Fig. 4E-G)**. On the other hand, the domain swap observed in design JH1 is correctly predicted by AF3 **(Extended Data Fig. 7B)** when prompted to fold it as a dimer. This is consistent with prior work which found that AF2 could be used to predict homo-oligomerization across the proteome^25^. Future design efforts might benefit from the inclusion of a filter to exclude such dimerization-prone designs.

The folding stability of water-soluble proteins is driven largely by the hydrophobic effect wherein nonpolar residues are hidden from water in the core and polar residues are oriented toward water at the surface^26^. We observed during our monomer design campaign that the cores of native proteins were more polar than those of ProteinMPNN-designed homologues **(Fig. 3D)**. De novo designed proteins are often hyperstable, while many native proteins are only marginally stable^4^. It is therefore reasonable to hypothesize that the increased polarity of native protein cores may be responsible for their decreased stability. Despite this expectation, seven of the monomer designs were stable above 95°C while possessing native-like core polarity **(Fig. 3C, Extended Data Fig. 6).** These findings suggest that elevated core polarity does not necessarily preclude hyperstability. This is consistent with recent rational design efforts incorporating buried charged residues into artificial coiled coils^27^. On the other hand, only one of the six most polar monomer designs JH1 was successfully characterized **(Fig. 3D)**, and it demonstrated an unexpected domain swap and two-stage melting curve **(Fig. 3C, top)**. However, most of the failed designs were those with β-sheet-rich folds, which are known to be more difficult to express than helix-rich designs^4,10^. It is therefore possible these designs may have failed for other reasons than their polarity, such as aggregation during folding.

While HBDesigner does not explicitly model water, it does implicitly account for it with some of its metrics, such as saturation and BUHs. Despite this, structural water intrusions into our designed networks occurred in two of the crystal structures we solved, those of monomer JH1 **(Fig. 3E, bottom)** and homodimer S2B **(Fig. 4E)**. This indicates that water can play an important role in network packing and plasticity, and that including explicit water in design may improve the accuracy of our design models. Encouragingly, recent work on GPCR redesign has suggested that solvent-mediated H-bonding interactions are both designable and important for allosteric signaling^28^. Still, our results also show that certain networks can tolerate modest rearrangements without losing stability or disrupting the overall fold. This is consistent with prior work on cold unfolding proteins, which observed similar plasticity in a designed partially hydrophilic core^29^.

Design of polar interfaces has been recognized as a longstanding challenge in protein design^30,31^. Based on this precedent, we expected that more polar interfaces would be more difficult to design. Indeed, we observed lower dimerization success rates for more polar homodimer designs, with a 33% success rate for the top half (5 of 15), compared to 81% (13 of 16) for the bottom half **(Fig. 4B)**. Several of our designs achieved similar interface polarity scores to those of the native obligate homodimers **(Fig. 4B)**, demonstrating that de novo designed oligomers can tolerate native-like interface polarity. Multiple other studies have used ProteinMPNN-based pipelines to improve polarity in designed interfaces^5,32^, although without direct comparison to native interfaces. Our method offers similar advantages but with more control over network properties and without reliance on evolutionary profiles.

Most of our dimer designs relied upon parallel or antiparallel helical packing commonly found in de novo designed assemblies^10,33^, while two designs used β-strand pairing to form either part (S2G) or all (S2H) of the interface **(Extended Data Fig. 9)**. This enabled elevated interface polarity (33% and 62% for S2G and S2H, respectively) despite relatively small sidechain H-bond networks. However, we observed significant oligomerization of both proteins by mass photometry, particularly S2H **(Extended Data Fig. 9).** Recent work on de novo binder design has highlighted β-strand pairing as a strategy for improving binder specificity toward certain targets^34^, although this relies on the target having exposed β-strands with non-canonical geometry to avoid aggregation. Another study has suggested that increased use of loop regions at the interface can facilitate more polar designs at the cost of more structural ambiguity^32^. Our approach serves as an orthogonal strategy for achieving polar interfaces primarily through sidechain-sidechain interactions, which is amenable to any backbone topology and therefore more generalizable.

Our heterodimer specificity experiment **(Fig. 5)** was motivated by the hypothesis that, for sufficiently similar scaffolds, H-bond networks could offer a way to impart specificity where shape complementarity and nonpolar packing may fail. Our results support this premise, as the HBDesigner-generated heterodimers displayed improved specificity despite sharing nearly identical predicted structures (TM score>0.95) and highly similar sequences (sequence identity>60%) with their nonspecific ProteinMPNN counterparts **(Fig. 5A)**. This is consistent with prior work using MC HBNet to design specificity into coiled coil assemblies^3,15,35^. We also observed reduced on-target affinity in the HBDes series **(Fig. 5C)**, which suggests that there may be a trade-off between affinity and specificity with respect to interface polarity. Prior work on nanoparticle design showed a similar trade-off between polarity and subunit assembly success rates in their heterodimeric interfaces^32^. Bioinformatic studies have also found that obligate homodimers typically possess more hydrophobic interfaces than transient ones^36,37^. Still, given that these designs were obtained from a one-shot design effort, it is likely that their affinity could be improved with more extensive iterative design techniques^38,39^.

The AF3 ipTM score and its derivatives are a commonly used proxy for binding likelihood, and they have been shown to be moderately discriminative between binders and nonbinders across many targets^35,38,40^. We therefore hypothesized that ipTM could be used to predict specificity by screening on-target and off-target A/B subunit pairs. Indeed, the ipTM scores correctly predict that the HBDes series will show reduced off-target binding compared to the MPNN series **(Fig. 5B-C)**. However, while the on-target ipTM is nearly identical for HBDes (0.90-0.91) and MPNN (0.91-0.93) designs, the on-target binding is 10- to 100-fold lower for most HBDes pairs. In this one limited example, AF3 ipTM served as a useful specificity screening tool but was unable to distinguish between weak and strong binders.

One limitation of our current approach is that it cannot be conditioned with specific atomic motifs, which would allow more fine-grained control of the designed network geometry. This would be useful for design of, for example, pH-sensitive networks, which require specific histidine-centered donor-acceptor configurations^13,41^. Our method also does not model ligands, which would enable scaffolding of ligand-binding proteins through intramolecular H-bonds^42^, although our preliminary experiments **(Extended Data Fig. 4)** found that second-shell interactions could still be recovered without explicitly modeling the ligand(s). These aspects represent interesting directions for future model development. It has been shown that small backbone movements can have large impacts on sidechain H-bonding^8^. As such, HBDesigner might benefit from allowing for backbone flexibility during sampling. Finally, HBDesigner uses idealized sidechain bond lengths and angles, which may limit its ability to sample realistic network geometry. Future work might consider employing an all-atom approach similar to that of the stochastic denoising model PLACER^24^.

This work includes only some of the possible applications for our design method. It could also be used to modify an existing design by grafting networks into an existing sequence, or to perform one-sided interface design (e.g., binder design). To facilitate use in these different design scenarios, networks can be generated with HBDesigner according to various criteria, including size, amino acid composition, and burial, which is difficult to achieve with generalist sequence design tools. Given its speed, efficacy, and easy integration into established protein design workflows, we expect that HBDesigner will be broadly useful for various applications. At the same time, we hope that our experimental design efforts have demonstrated that design of buried H-bond networks is tractable with modern protein design tools, opening new avenues for more ambitious design objectives.

## Methods

### Dataset curation

For model training and validation, we curated a modified version of the ProteinMPNN dataset^16^, which is a snapshot of the Protein Data Bank from August 2021 containing only protein chains with all ligands, nucleotides, solvent, and cofactor atoms removed. Biological assemblies solved by x-ray crystallography or cryo-EM with resolution ≤ 3.5 Å and 10-10,000 total residues with ≥25% of residues resolved were collected. Sidechains were decomposed into chi angles and rebuilt using idealized bond lengths and angles. PyRosetta^19^ was used to add and optimize polar hydrogens. Sidechain-sidechain H-bonds were calculated using the Rosetta REF2015 energy function^43^ and used as edges to construct an H-bond graph for each assembly **(Fig. 1B)**. H-bond networks were identified using connected-component analysis, and those with 2-6 residues were retained. To avoid transient surface networks, we kept only networks for which ≥50% of residues were buried (>4.0 weighted sidechain neighbors as determined by the PyRosetta LayerSelector). The curated assemblies were split chain-wise into sequence clusters (30% sequence identity) following the splits from ProteinMPNN. This resulted in 19,916/1,209/1,247 clusters assigned to the training/validation/test sets, respectively, containing 84,388/4,003/3,872 chains with 1,767,365/98,078/84,019 H-bond networks.

### Training procedure

During training, we cycle through a shuffled list of sequence clusters and sample a random assembly from each cluster, then sample a random H-bond network for each assembly. Each assembly-network pair constitutes one training example. The sequence and sidechain coordinates of all non-network residues are cleared prior to featurization. We train both the design and packing model using PyTorch^44^ for 200,000 optimizer steps using a batch size of 10,000 tokens. We use the Adam optimizer with the Noam learning rate schedule^45^ with a warmup period of 4,000 steps followed by an exponential decay for the remainder of training.

### Design model architecture

The design model **(Fig. 1A, top)** is implemented as a message passing neural network (MPNN) in which protein residues are represented as nodes in the molecular graph, with edges connecting residues to their 30 nearest neighbors (assigned based on Cβ-Cβ distances). Geometric information derived from the backbone structure is encoded as a combination of node- and edge-level features. Conditioning information influences the model at each layer through conditional features. See **Supplementary Table 1** for a summary of all input features.

The architecture of the design model is summarized in **Extended Data Fig. 11A**. The initial node features for each residue consist of backbone dihedrals and a partially masked amino acid sequence. To generate initial edges between residues, pairwise distances between neighboring backbone atoms (N, C, O, Cα, and virtual Cβ) are each encoded with 16 radial basis functions (RBFs) evenly spaced from 0 Å to 20 Å. The other edge feature is an encoding of relative sequence (or chain) separation of neighbors within ± 32 residues. The node and edge features are then updated with three sequential message passing layers which propagate node and edge updates throughout the molecular graph. These layers use a similar message passing scheme to the decoder layers from ProteinMPNN. Prior to each message passing step, vectors encoding the various forms of conditional information are used to update the nodes using a simple node conditioning block, where is the node embedding and is the conditioning vector:

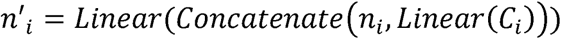

Two output predictions are calculated from the final node representations: 1) a binary prediction of whether each residue is in the network, and 2) a categorical prediction of the amino acid identity of each network residue.

### Design model training

To avoid overfitting, we apply random Gaussian noise to the backbone coordinates with a standard deviation of 0.2 Å, as introduced by ProteinMPNN. For each training example of a network with N residues, a decoding step is randomly sampled such that T residues are masked and must be predicted, where *T*∊ {1, …, *N*}. To save memory, only protein chains containing one or more network residues are provided to the model during training. The design model was trained using a weighted sum of positional and sequence loss terms:

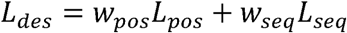

Where *L_pos_* is a binary negative log likelihood loss:

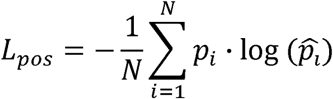

Where *p_i_* is the ground-truth probability and 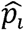 is the predicted probability of residue position *i* to participate in an H-bond network of N residues. We define *p_i_* as a uniform distribution across the masked residue positions with all other positions set to zero. We formulate *L_seq_* as a multi-class focal loss^46^:

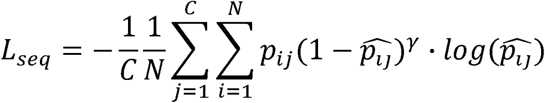

Where *p_i_* is the ground-truth probability (binary label) and 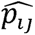 is the predicted probability of amino acid *j* at position *i*, where C is the number of valid amino acid types. For our production models, we set *w_pos_* = *w_seq_* = 1 and *γ* = 2 based on results of hyperparameter tuning. The design model includes 1.27M trainable parameters and can be trained in 10 hours on an A100 GPU.

### Design model conditioning

We sought to train one general-purpose design model capable of supporting both conditional and unconditional generation. To do this, we randomly provide or mask each type of conditioning information for each training example. Virtual guide atom features are sampled 50% of the time, while full and partial sequence conditioning are each sampled 10% of the time. The guide atom and sequence conditioning features are sampled independently, so both or neither may be provided for some examples.

Sequence conditioning **(Fig. 1C)** enables residue-level control over the amino acid composition of the network. This information is encoded as a vector of 20 weights, one for each amino acid, expressing the amino acid composition of the network. When encoding the full sequence of a network, this would be expressed as 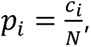, where *c_i_* is the number of residues with amino acid type *i* for a network of size *N*, and 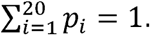. This formulation also allows us to encode partial sequence conditioning for networks where the amino acid type of one or more residues may be unknown. To encode an unknown residue, we simply spread the probability for that residue equally across all valid polar amino acids. This scheme also supports ambiguous (e.g., “either GLU or ASP”) sequence conditioning. During training, sequence conditioning is a soft constraint used to bias predicted position and sequence logits. During inference, it is enforced as a hard constraint, so that only amino acids included in the options for sequence conditioning are permitted. Anchor residue conditioning **(Fig. 1E)** also acts as a hard constraint, such that all provided anchor residues are guaranteed to be included in the predicted networks.

The virtual guide atom **(Fig. 1D)** encodes an approximate center-of-mass for the network in 3-dimensional space, inspired by the ORI token from RFdiffusion2^22^. During training, this virtual atom is generated by calculating the centroid of the Cβ atoms of the network, then sampling a point from a normal distribution located at the centroid with a standard deviation of 4 Å. We then calculate pairwise distances between the virtual atom and all Cβ atoms in the protein and encode them with RBFs, as previously described. By default, guide atom conditioning is provided as a soft constraint during inference, but it can be converted to a hard constraint by specifying a maximum radius around the guide atom which is used to set designable residues.

### Packing model architecture

The HBDesigner packing model **(Fig. 1A, middle)** uses a similar architecture to the design model, with a few changes **(Extended Data Fig. 11B)**. Most notably, it does not accept conditional information, since the network residues have already been assigned, so each decoder block includes only a message passing step. The packing model also uses three rounds of recycling, wherein the predicted sidechains are recycled back through the same model to further refine predictions^17^. We introduce an additional node feature to encode the sine and cosine of these predicted sidechain dihedrals. Pairwise distance RBFs for these sidechain heavy atoms with respect to all neighboring atoms are also added to the edge features. All chi angles are zeroed out and used to rebuild the sidechains prior to the first pass through the model. The final node representations are used to predict the sine and cosine of the chi angles, and sidechains are then constructed using idealized bond lengths and angles.

### Packing model training

Similar to the design model, we apply random Gaussian noise to the backbone coordinates, in this case with a standard deviation of 0.02 Å. To save memory, input proteins are cropped to include only residues within 10 Å of the network centroid (calculated with Cβ atoms). Unlike the design model, amino acid types of all network residues are provided unmasked for every training example. During training, the number of recycles *r* for each example is uniformly sampled, *r* ∊ {0,1,2,3}. The packing model was trained on a weighted sum of multiple loss terms:

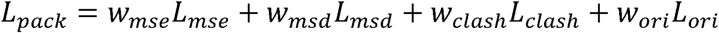

Where *L_mse_* is the mean-squared-error of the predicted chi angle sine 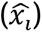 and cosine 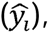, reweighted by chi angle abundance factor *M_χi_*, with an added normalization penalty *L_form_*:

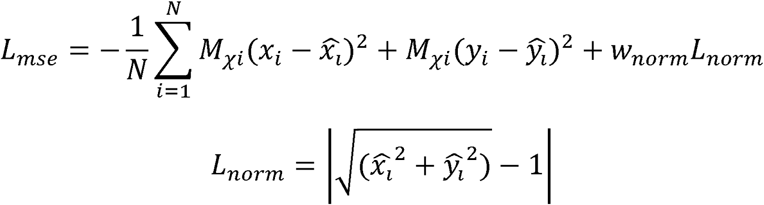

The chi abundance factor is obtained by calculating the number of chi angles 1-4 present in each batch (*N_χi_*), then converting this to a relative weight (*M_χi_*) such that rare The chi abundance factor is obtained by calculating the number of chi angles 1-4 chi angles receive larger weights:

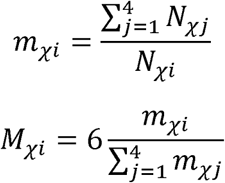

*L_msd_* is the mean-squared-deviation of the predicted sidechain coordinates 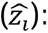:

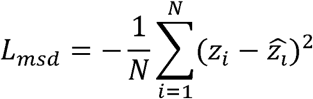

*L_clash_* is a clash loss adapted from OpenFold^47^ penalizing overlap of van der Waals radii of predicted sidechain atoms with other backbone and sidechain atoms:

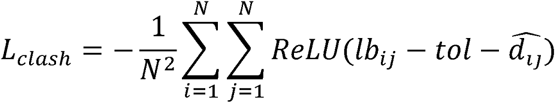

Where *lb_ij_* is the allowed lower bound of interatomic distances based on vdW radii, *tol* is an overlap tolerance (set to 1.5 Å), and 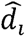 is the predicted interatomic distance between atoms *i* and *j*. *L_ori_* is an orientation loss calculated on the mean-squared-deviation of the pairwise distances between all predicted sidechain atoms in each network:

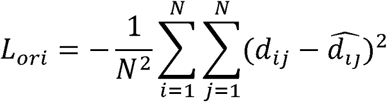

For our production model, we use *w_mse_* = *w_msd_* = 1.0, *w_clash_* = *w_orient_* = 0.2, and 0.1. The packing model includes 1.21M trainable parameters and can be trained in ∼40 hours on an A100 GPU.

### Scoring module implementation

HBDesigner uses PyRosetta to minimize and score the predicted networks **(Fig. 1A, bottom)**. The MinMover class is used with a tolerance of 1e-6 and maximum of 1,000 *lbfgs_armijo_nonmonotone* minimizer iterations. Cartesian minimization is enabled using the full REF2015 score function, and all atoms are fixed except the sidechain atoms of the network residues. Minimized networks are first checked for validity, meaning that all network residues are participating in a single contiguous network of sidechain-sidechain H-bonds. If valid, the network is scored with the metrics described in the MC HBNet paper^8^, which are intended to reward highly connected, energetically favorable networks. These metrics, which capture contributions from both sidechain-sidechain and sidechain-backbone H-bonds, include buried unsatisfied heavy polar atoms (BUHs) and buried unsatisfied polar hydrogens (BUPHs), saturation, and the HBNet Score. Saturation estimates the fraction of potential H-bonding donor/acceptor sites being satisfied by H-bonds in a set of residues for a given geometry. HBNet Score is the difference in energy between the minimized network and an equivalent backbone with all residues replaced by glycine.

### HBDesigner inference

Once trained, HBDesigner can be applied to new proteins in inference mode. Given an input backbone and, if applicable, conditioning information, the design model is repeatedly sampled to collect many unique potential networks (typically 100-1,000 per backbone). In sampling mode, the design model autoregressively assigns network residues until the desired network size has been reached. At each decoding step, the position of the next network residue is predicted, then its amino acid identity is assigned. On subsequent steps, these decoded residues are used to update the node embeddings which are fed back into the model, now representing a partially decoded network. Once fully sampled, the backbones are cropped, and their network sidechains are predicted with the HBDesigner packing model. They are then minimized and scored with various H-bonding metrics using PyRosetta, as described above. Any predictions that fail to form contiguous networks of H-bonds are discarded, and additional filters (e.g., saturation) can be used to further trim the set of returned networks. By default, we follow the MC HBNet convention and return only networks with ≤0 BUHs, ≤5 BUPHs, ≥0.5 saturation, and ≤0 REU HBNet Score. Networks are then sorted by these same metrics, prioritized in the order written, and the top K networks are returned. As shown in **Table 1**, total runtime for HBDesigner is under 10 seconds per scaffold for most common use cases.

### Designed network native likeness test

To evaluate native likeness of our designed networks **(Fig. 2A, Extended Data Fig. 1)**, we iterated over the clusters of our test dataset and randomly sampled native networks (n=1,000 networks unless otherwise stated) to obtain a baseline. We then repeated this sampling with the same random seed but allowed HBDesigner to instead design its own network onto the native backbone and compared the resulting metrics. Amino acid abundance and pairwise residue enrichment **(Fig. 2A, Extended Data Fig. 1A)** were calculated on all sampled networks, while packing metrics **(Extended Data Fig. 1B)** were collected only for designs which were successfully packed into a valid contiguous network. Model conditioning success rates **(Extended Data Fig. 2)** were evaluated using the same sampling loop, but with each conditioning channel (sequence, virtual guide atom, anchor residue) toggled on or off to assess the impact on native recovery metrics.

### Native network repacking test

To assess the efficacy of different packing methods, we tested their ability to repack native networks on their native backbones without additional sequence context **(Fig. 2B)**. We calculated rotamer recovery by considering any prediction within ± 10° of the native sidechain as successful. The Rosetta packing method used the PackRotamers mover with only H-bond (hbond_sc) and repulsive (fa_rep) terms enabled with -ex1 -ex2 -ex3 rotamer sampling, which gave the best performance at a reasonable runtime. The PIPPack comparison was done using 3 recycles and an ensemble of 3 models with no postprocessing.

### Refolding benchmark

We evaluated HBDesigner and comparable methods using the refolding test outlined in **Fig. 2C**, applied to three different sets of scaffolds. We obtained 532 native monomeric domains of 50-200 residues from the HBDesigner test set by randomly sampling one representative from each qualifying sequence cluster. We also generated 726 de novo monomeric domains of 80-120 residues using RFdiffusion^48^ with fold conditioning based on secondary structures from native monomers of the same size from the test set clusters. Finally, we generated 690 de novo heterodimers of 200-300 total residues using unconditional generation with RFdiffusion. The de novo generated scaffolds were filtered for suitable secondary structure using STRIDE^49^ to ensure ≥50% helix/sheet content and ≥10% turn/coil content. The de novo monomers were further filtered by checking that ≥12% of the residues were classified as Core by the Rosetta LayerSelector to filter out non-globular backbones and ensure a well-defined core for network design. The de novo heterodimers were filtered with the Rosetta InterfaceAnalyzer^30^ to include only pairs with 500Å δ ΔSASA δ 2000Å, to avoid tangential interfaces and entwined topologies.

We ran HBDesigner in unconditional sampling mode on each set of scaffolds and collected the top 5 networks each for network sizes of 2-6 residues. Sampling parameters for HBDesigner are outlined in **Supplementary Table 2**. Large networks required more samples to find sufficient high-quality networks, at the cost of slightly increased runtime. We also utilized a low-noise version of the design model for larger networks based on better results in empirical testing. All polar amino acids were allowed, and at least one core residue was required in each network to ensure sufficient burial.

We used LigandMPNN^50^ to generate 4 sequences per network, provided the HBDesigner network as both sequence and structural context (using the *-- ligandmpnn_use_side_chain_context* flag). De novo monomer and heterodimer runs used sampling temperatures of 0.2 and 0.1, respectively, sampled with the LigandMPNN model trained with 0.1 Å backbone noise. For the native monomers, we re-trained ProteinMPNN^16^ using the reported training splits with 0.2 Å noise to avoid data leakage, since the default checkpoint is trained on all sequence clusters. We then refolded all sequences using a high-throughput AlphaFold3^20^ procedure using no MSAs or templates, 1 model seed, 1 diffusion sample, and 1 recycle. All successfully refolded sequences (pLDDT ≥ 80 or ipTM ≥ 0.7 and RMSD_Cα_ ≤ 2.0 Å) were then minimized with PyRosetta, and buried H-bonds were calculated for each design. For the scaffold recovery test **(Fig. 2G)**, we consider a scaffold successfully recovered if any of the 20 designed sequences (4 per network) is well-folded (see above) and also recovers all designed network H-bonds with an RMSD_net_ ≤ 1.5 Å.

### Comparison with literature methods

To compare with baseline ProteinMPNN **(Fig. 2D-E)**, we used the settings outlined above to generate 4 full sequences for each scaffold. To implement a bias toward buried polar amino acids in ProteinMPNN, we add a constant bias favoring all amino acids capable of sidechain H-bonding (DEHKNQRSTWY) to all buried (or interface) positions. We tuned the value of this bias term to align the fraction of buried polar residues with those produced by HBDesigner for each scaffold set. This resulted in bias terms of 1.5, 1.0, and 0.5 for native monomers, de novo monomers, and de novo heterodimers, respectively. The Rosetta MC HBNet protocol^8^ was run in PyRosetta using settings configured to match HBDesigner as closely as possible, including network size, minimum burial, number of core residues, and ranking criteria. Rotamer sampling levels -ex1 -ex2 were used, and 10,000 Monte Carlo samples were drawn for each scaffold. Interface design used the HBNetStapleInterface class to ensure cross-chain networks.

### Second-shell interaction recovery benchmark

To assess the ability of HBDesigner to recover second-shell H-bonding interactions, we collected two datasets of native protein-ligand binding sites. The first was curated from the AME dataset introduced by RFDiffusion2^22^, which includes 41 diverse enzyme active sites with curated theozyme definitions. We used PyRosetta to identify H-bond networks of 2-6 residues in each structure terminating at one of the theozyme residue sidechains. If more than 50% of the network was comprised of theozyme residues, we randomly downsampled to keep only 50% as anchor residues. This produced 54 networks with 96 designable (non-anchor) residues spread across all 41 PDBs. The second dataset was curated of a subset of sequence clusters from the MetalPDB^21^, which is a database of metal-binding sites in naturally occurring proteins. We used a similar procedure to identify H-bond networks terminating at one or more metal-binding residues, downsampling as necessary. This produced 67 sites with 112 designable residues spread across 50 PDBs. HBDesigner was run on each site with 500 samples and the native anchor residues provided as conditioning, and the top-10 ranked designs were kept. Non-anchor residues present in one or more of the predicted networks were considered recovered.

### Computational binding specificity test

To assess the impact of HBDesigner on binding specificity **(Fig. 2H)**, we first obtained a set of designable de novo heterodimer scaffolds by generating backbones with RFdiffusion, then designing sequences with ProteinMPNN and refolding them with AlphaFold3. From those heterodimers with high confidence values (ipTM > 0.8, pTM > 0.7), we collected 19 structures with diverse interface geometries. These were used as seeds for RFdiffusion partial diffusion, where we used *partial_T* = 5 with default settings to generate 10 highly similar backbones from each seed, producing 190 total scaffolds. We then designed a single full sequence for each backbone using the LigandMPNN model *v_32_010_25*, omitting cysteine residues, and using a temperature of 0.1. When using HBDesigner, we generated networks of 4 residues with a minimum burial of 3.0 weighted sidechain neighbors and up to 2,000 samples per backbone, keeping the top 30 networks found. LigandMPNN was then used to design the remaining sequence with the same settings as above, except with the network residues provided as context. Within each backbone family, we selected the top-ranked non-redundant network for each backbone, meaning that no two family members had more than 2 shared network positions. We then assembled all possible pairs of A/B subunit sequences (190 for correct pairing, 1,710 for mispairing) and predicted each heterodimer using AlphaFold3, as described above. To assess interface confidence, we use the interface predicted template modeling (ipTM) score, which has been shown to correlate with DockQ score and other measures of interface quality^51^.

### Computational design of de novo monomers

We obtained de novo monomer backbones of 80-120 residues using RFdiffusion, generating 1,000 backbones each with four different settings: 1) unconditional generation, 2) fold-conditioning on native monomer folds, 3) using the *monomer_ROG* potential, 4) using *monomer_contact* potential. Backbones were filtered as described above. HBDesigner was used to design 3-4 residue networks with unconditional sampling, with a minimum burial of 4.0 weighted sidechain neighbors and at least 1 network residue in the core. The top 5 networks for each backbone were obtained, and LigandMPNN was used as described above to sample 4 sequences per network. This produced a total of 16,202 H-bond networks with 64,808 unique sequences, which were then refolded with AlphaFold3 and filtered with various metrics.

First, we filtered for highly confident, self-consistent backbones (pLDDT ≥ 80, RMSD_Cα_ ≤ 1.0 Å), which left 30,572 designs. Next, we minimized each design with PyRosetta and checked whether all HBDesigner-predicted H-bonds were recovered in the refolded structure. We also removed any designs with RMSD_net_ > 1.0 Å. This left 3,245 high-quality designs, which we filtered further by BUHs (<5) and core polarity (>28%). Finally, we analyzed the remaining 544 designs with PLACER, a model trained to predict sidechain flexibility^24^. We generated an ensemble of 50 samples for each design and filtered for high H-bond recovery (≥ 80%) and low RMSD_net_ (≤ 0.5 Å), leaving a final set of 271 designs. From these, we selected 19 designs for expression and characterization based on visual inspection of network quality and centrality, as well as structural diversity.

To redesign loop 1 for protein JH1 to avoid the dimerization involving domain swapping, we used the motif scaffolding protocol from RFDiffusion^48^ to generate alternative loop conformations of varying length (3-12 residues). We then used ProteinMPNN^16^ to design sequences for the loop and AlphaFold2^52^ to predict structures of each design as both a monomer and a dimer. We scored all sequences based on high confidence (mean pLDDT) of monomer prediction and low confidence of dimer prediction. We selected the top scoring sequences and used AlphaFold3 prediction to choose the sequence with the lowest ipTM value for the dimer interface.

### Calculation of core and interface polarity metrics

The core polarity metric shown in **Fig. 2D** was calculated by counting the polar amino acids (DEHKNQRSTWY) found in the core and dividing them by the total number of residues in the core. Only those designs that refolded well (pLDDT ≥ 80, RMSD_Cα_ ≤ 2.0) with ≥10 core residues were included to avoid counting disordered proteins. For interface polarity **(Extended Data Fig. 3B)**, we used the InterGroupInterfaceByVectorSelector to select interface residues. Only designs with ipTM ≥ 0.7 and RMSD_Cα_ ≤ 2.0 and ≥20 interface residues were included. Buried core H-bonds **(Fig. 2E, Extended Data Fig. 3A)** were calculated as all sidechain-sidechain or sidechain-backbone H-bonds including at least one core residue after minimization. Buried interface H-bonds **(Extended Data Fig. 3B)** had the added constraint that each bond had to connect residues in two different chains.

When assessing the core polarity of our tested designs **(Fig. 3C)**, we obtained more informative results using a different metric, percent buried polar surface area (bSA%), which we formulated to capture the contribution of each individual atom rather than each residue. To calculate this, we used PyRosetta to select all core and boundary residues (SASA < 40). We then calculated the SASA for each residue and subtracted it from the theoretical maximum to get an estimated per-residue absolute buried surface area (bSA). We then calculated the bSA for an equivalent poly-glycine backbone and subtracted this value for each residue to remove backbone contributions. We repeated this calculation, including only polar atoms, and divided polar bSA by total bSA to get bSA%. For interface polarity **(Fig. 4C)**, we use the InterfaceAnalyzer to calculate polar and total dSASA, then divide these to get percent polar dSASA, which includes both sidechain and backbone contributions. To compare our designs to native homodimers, we obtained a curated dataset of experimentally validated obligate homodimers (n=122) from ref ^25^. Since our designs were nearly all helical, we selected only those native homodimers with primarily helical interfaces (≥50% of interface residues assigned helical by STRIDE^47^). This left us with 47 validated native helical homodimers for comparison.

### Computational structural validation

To probe the quality of our designed structures, we ran several different, independently trained structure prediction models on our biochemically validated, well-expressing monomers (n=11) and homodimers (n=18) **(Extended Data Fig. 8)**. We used Boltz-2^53^, ESMFold2^54^, and Chai-1^55^, and we include AF3, which we used during design, for reference. None of the other three tools were used to select designs for expression. All folding models were used in single-sequence mode (no MSAs or templates) with 1 diffusion seed and 1 sample and the default number of diffusion steps and recycles/loops for each method, except ESMFold2 which used n_loops=3 and 50 steps. Alignment metrics were calculated against the design models obtained from AF3 folding and PyRosetta energy minimization in our original design campaigns.

We also used PLACER^24^ to assess the packing quality of our designed networks. To do this, we generated n=50 PLACER samples for each design, using a randomly selected network residue as the *target_res* which defines the center of the predicted crop. For the homodimers, used only the proximal copy of the network for evaluation, since the second copy was often omitted from the crop. Network sidechain RMSD was calculated against the original design model described above.

### Computational design of de novo homodimers

We generated de novo homodimer backbones of 60-120 residues by RFdiffusion using C2 symmetry and auxiliary potential *olig_contacts*, with intrachain and interchain weights set to 0.4 and 0.2, respectively. The resulting scaffolds were filtered by radius of gyration to ensure backbones were compact, and any designs with entwined topologies were removed. We ran HBDesigner to design two-, three-, and four-residue networks across the interface, collecting the top 15 networks per scaffold. We used symmetry constraints (flag *--symm_chains* A,B) to select only networks which could be symmetrized across the interface without introducing clashes. This means that, for example, each three-residue network design actually has two copies of that network including six total residues. All polar residues were allowed, with a minimal burial value of 3.0 weighted sidechain neighbors in the interface.

To design the remaining sequence, LigandMPNN was used to generate 5 sequences per network, providing the HBDesigner network as sequence context. We used a sampling temperature of 0.1 and *--homo_oligomer* flag turned on to automatically set symmetric residues and equal symmetry weights. We then refolded all sequences using AlphaFold3, as previously described. All successfully refolded sequences (average pLDDT ≥ 80, ipTM ≥ 0.8, RMSD_Cα_ ≤ 2 Å) were minimized with PyRosetta and filtered based on full recovery of the designed H-bond network.

Multi-network designs were generated by collecting all single-network designs that passed all design filters and attempting to graft in an additional network sampled for the same backbone. To avoid steric clashes, combined networks were rejected if any sidechain heavy atoms from individual networks were separated by less than 3 Å, or if any positions would be occupied by both networks at once. An additional round of sequence design, refolding, and filtering was then performed as described above.

### Protein expression and purification

DNA sequences were ordered as gene fragments from Twist Bioscience and inserted into a Pet28(b) vector for expression containing a N-terminal His-tag and PreScission protease cleavage site. Plasmid vector for homodimer proteins contained a N-terminal His-tag, Maltose-binding protein (MBP), and PreScission protease cleavage site. Heterodimer plasmid vector contained a N-terminal His-tag and AviTag or FLAG-tag. DNA constructs were transformed into BL21(DE3) Escherichia coli cells with 50 ng DNA and plated onto LB agar plates supplemented with 50 µg/mL Kanamycin selection antibiotic and incubated overnight at 37°C. Transformed colonies were picked and inoculated into 5 mL LB supplemented with 50 µg/mL Kanamycin selection antibiotic overnight at 37°C with shaking at 225 rpm. For small scale expression, 2 mL of overnight culture was added to 100 mL fresh TB supplemented with 50 µg/mL Kanamycin selection antibiotic and 800 µL of 50% glycerol. For large scale expression, 15-20 mL of overnight culture was added to 1 L fresh TB supplemented with 50 µg/mL Kanamycin selection antibiotic and 8 mL of 50% glycerol. Cells were cultured at 37°C for 2-3 hours or until an optical density of 600 nm of 0.7-0.8 was measured. Protein expression was induced by addition of Isopropyl β-D-1-thiogalactopyranoside (IPTG) at a final concentration of 750 µM, followed by incubation overnight at 16°C with shaking at 225 rpm. Cells were pelleted by centrifugation and stored at -20°C.

Cell pellets were thawed at room temperature and lysed with Lysis Buffer (50 mM Tris, 500 mM NaCl, 10 mM Imidazole, pH 8.0) supplemented with 1X PMSF, 1X Bestatin, 1X Leupeptin, 1X Pepstatin, 1X B-PER, 1 mg/mL Lysozyme, and 0.145 µL/ 1 mL Benzonase by shaking 1-2 hrs at 25°C. Lysate was centrifuged at 15,000 rpm for 30 min and passed through a 5 µm Titan3 syringe filter (Thermo Fisher, 45025-NN). Expressed protein was purified by Nickel purification. For purification of proteins from small scale production, briefly, batch binding of filtered lysate was performed by incubating with Ni-Penta Agarose 6 (Marvelgent Biosciences, 11-0228) at 4°C for 1-2 hrs with gentle rocking. Resin was collected in 24-well Whatman Uniplate filter plate (Whatman, 7700-9904) and centrifuged for 1 min at 25°C at 250 × g. Resin was then washed sequentially with High Salt Buffer (50 mM Tris, 1 M NaCl, 25 mM Imidazole, pH 8.0) and 1X PBS (pH 7.4) by centrifugation for 1 min at 25°C at 250 × g. Protein was eluted by Elution Buffer (1X PBS, 500 mM Imidazole, pH 7.4) and centrifugation for 1 min at 25°C at 250 × g. Eluted protein was concentrated via 10 kDa MWCO 0.5 mL Amicon concentrator (Millipore Sigma, UFC501096) and centrifugation at 14,000 × g. Concentrated protein was buffered exchanged into PBS via 0.5 mL 7k MWCO Zeba Spin Desalting Column (Thermo Fisher, 89882) following the manufacturer’s protocol. In brief, for purification of proteins from large scale production, batch binding of filtered lysate was performed by incubating with Ni-Penta Agarose 6 (Marvelgent Biosciences, 11-0228) at 4°C for 1-2 hrs with gentle rocking. Resin was collected in poly-prep columns and washed with 10 bed volumes (BV) High Salt Buffer and 10 BV 1X PBS, and eluted with Elution Buffer.

For JH1 crystallization experiments, PreScission protease was added to purified protein at a 1:100 mg ratio, then transferred to a 1 kDa dialysis bag and dialyzed against 1xPBS 5 mM BME buffer. For homodimer crystallization experiments, PreScission protease was added to purified protein at a 1:50-1:250 mg ratio in Protease Cleavage Buffer (50 mM Tris HCl, 150 mM NaCl, 1 mM 2-Mercaptoethanol, pH 7) and rocked overnight at 4°C. Nickel purification was used to remove cleaved tag and uncleaved protein, as previously described. GST-tagged PreScission protease was removed by gluthathione resin.

### Protein crystallography and model refinement

Following protease cleavage, proteins were purified by gel filtration via S75 Superdex column in Crystallization Buffer (20 mM Tris, 100 mM NaCl, pH 7.4) at 4°C and concentrated as described above. Using the sitting drop vapor diffusion method, crystallization screens were set up via Mosquito liquid handler (SPT Labtech) with drops consisting of 150 µL protein and 150 µL reservoir solution. Crystal screens were placed at 20°C and checked regularly. JH1 protein crystals were obtained by crushing the initial crystal solution formed using SG-1 (Molecular Dimensions) screening solution and seeding in the same mother solution (20% w/v PEG 3350, 0.2 M MgCl2*6H2O). Protein crystals were cryoprotected for x-ray diffraction by sequential mixing with 6.25%, 12.5%, and 25% polyethylene glycol, prepared from serial dilution of polyethylene glycol and reservoir diluent at a 1:1 ratio, then flash frozen by submersion in liquid nitrogen.

Protein crystal x-ray diffraction data was collected at SouthEast Regional Collaborative Access Team (SER CAT) with the Beamline 22-ID-D synchrotron. Diffraction data was reduced using PROTEUM5 (Bruker, 2025.6-0) and the solution was solved by Molecular Replacement in PHASER-MR by using the AlphaFold3 prediction as the starting model. Iterative model refinement was performed using Phenix (2.0.5936) and model building was done with Coot (WinCoot 0.9.8.92). Data collection and refinement statistics are provided in **Supplementary Table 3**.

### Circular dichroism (CD)

All protein samples were diluted to 20 μM in 1xPBS. The measurements were obtained using a JASCO-1500 instrument in a 1 mm cuvette. The CD spectra was measured from 260 nm to 190 nm at 25°C. The melting temperature curve was obtained by measuring CD signal at 222 nm from 5°C to 95°C in 1C steps.

### Homodimer mass photometry

Mass photometry was performed on a Refeyn OneMP instrument. Protein solutions were prepared at a concentration of 100 nM in PBS immediately prior to measurement. Droplet-dilution was performed by addition of equivalent volume to PBS on a glass coverslip. Videos were collected for 60 s and analyzed by DiscoverMP software (Reyfen) to determine the number of binding events. Protein size was calculated using Thermo Fisher NativeMark Protein Standard (Thermo Fisher, LC0725).

### Heterodimer computational specificity screening

To identify families of de novo heterodimers for expression, we reexamined the results of the computational specificity test as follows. Within each family, we first selected backbones for which both the ProteinMPNN and HBDesigner sequences had high on-target ipTMs (≥0.75). We then cross-referenced these hits with one another, looking for pairs of backbones for which the off-target ipTMs were high (≥0.75) for the ProteinMPNN sequences, but low (≤0.50) for the HBDesigner sequences. The top 5 cliques of compatible backbones were then further screened by running each subunit through AlphaFold3 as a monomer and as a homodimer. Any subunit with monomer pLDDT < 80 or homodimer ipTM > 0.5 was penalized, and the clique with the fewest penalties was selected for expression.

### Heterodimer biotinylation and ELISA binding assay

Purified protein was buffer exchanged into 10 mM Tris at pH 8.1. For biotinylation, 40 µM of protein was mixed with BiomixA, BiomixB, and BirA, brought to a final volume of 135 µL with DI H2O, and incubated at room temperature. To remove reaction components, the mixture was repeatedly concentrated via 10 kDa MWCO 0.5 mL Amicon concentrator and diluted with buffer.

ELISA plates were prepared by coating 96-well MaxiSorp plates (Thermo Fisher, 439454) with 50 µL of 1 µg/mL Neutravidin overnight at 4°C, then washed with 0.05% Tween-20 in phosphate buffered saline (PBST) the following day. Blocking was performed with 1% casein for 1 hr at room temperature with gently shaking. Plates were then treated with 100 nM of biotinylated protein for 1.5 hrs with gently shaking and thoroughly washed with PBST. Plates were subsequently treated with protein in PBST with 0.1% casein for 45 minutes, rinsed, and incubated with a 1:5,000 dilution of HRP anti-DYKDDDDK antibody (Biolegend, 637311) in PBST with 0.01% casein for 30 min. Plates were then rinsed with PBST, and 50 µL per well of TMB substrate (Biolegend, 421101) was added and incubated for approximately 20 minutes. The reaction was stopped by addition of 50 µL of 1 N HCL, and absorbance was immediately measured at 450 nm with a reference wavelength of 570 nm.

## Supporting information

Design Models

Design Sequences

## Data and Software Availability

The code and model weights for HBDesigner are available on GitHub (https://github.com/RosettaCommons/HBDesigner). The preprocessed dataset used for training and validation is available on Zenodo^56^, and additional source data and evaluation scripts are provided on Zenodo as well^57^. The sequences and design models of all expressed proteins are provided as supplementary data. Crystal structures of the designed proteins have been deposited in the Protein Data Bank under accession codes 13JM, 13FC, 35SL, and 35SB.

## Acknowledgements

This work was supported by NIH grant R35GM131923 (B.K.), NSF grant MCB-2319819 (B.K.), and NSF fellowship DGE-2040435 (H.D., N.Z.R.), and by a Pre-doctoral Fellowship from the American Foundation for Pharmaceutical Education (H.D.). This work utilized the resources of the UNC Longleaf computing cluster. The authors would like to thank Rachel Clune and Hope Woods from the Rosetta Commons for their assistance in maintaining HBDesigner. Protein crystallization and data collection were performed in the UNC Protein Expression and Purification & Macromolecular Crystallography core lab supported by National Cancer Institute of the National Institutes of Health under award number **P30CA016086.** We would also like to acknowledge Dr. John Sondek’s lab, particularly Dr. Stuart Endo-Streeter for the technical assistance in data reduction, and Matthew Hvasta from Dr. Brian Kuhlman’s lab, for the technical assistance in molecular replacement.

## Author contributions

H.D., B.T.H., T.M., and B.K. conceptualized the study. H.D. and N.Z.R. wrote the software, and H.D. performed the computational experiments. J.T.H., H.D., and T.M. designed the monomers, and J.T.H. and T.M. characterized the monomers. T.M. and N.I.N. solved the JH1 crystal structure. B.T.H. designed and characterized the homodimers and solved the homodimer crystal structures with assistance from N.I.N. H.D. and B.T.H. designed the heterodimers, and B.T.H. performed the heterodimer binding experiments. H.D., B.T.H., and T.M. made the figures and wrote the manuscript, and all authors contributed to review and editing. B.K. supervised the project.

