## Supplementary Materials for "Deep learning based design of buried hydrogen bond networks with HBDesigner"

Table of Contents:

Extended Data Figures 1-11

**Extended Data Figure 1: Native likeness of designed hydrogen bond networks.**

**Extended Data Figure 2: Model conditioning success rates.**

**Extended Data Figure 3: Additional refolding benchmark results.**

**Extended Data Figure 4: Second-shell network design benchmark.**

**Extended Data Figure 5: Family-level analysis of designed heterodimer specificity.**

**Extended Data Figure 6: Characterization of additional monomers with highly polar cores.**

**Extended Data Figure 7: Additional characterization of design JH1.**

**Extended Data Figure 8: Computational structural validation.**

**Extended Data Figure 9: Characterization of additional homodimers with single interface hydrogen bond networks.**

**Extended Data Figure 10: Characterization of additional homodimers with multiple interface hydrogen bond networks.**

**Extended Data Figure 11: Architecture of the HBDesigner design and packing models.**

Supplementary Tables 1-3

**Supplementary Table 1: Description of input features for HBDesigner sequence design model.**

**Supplementary Table 2: HBDesigner inference settings used for the refolding benchmark.**

**Supplementary Table 3. Crystal data collection and refinement statistics.**

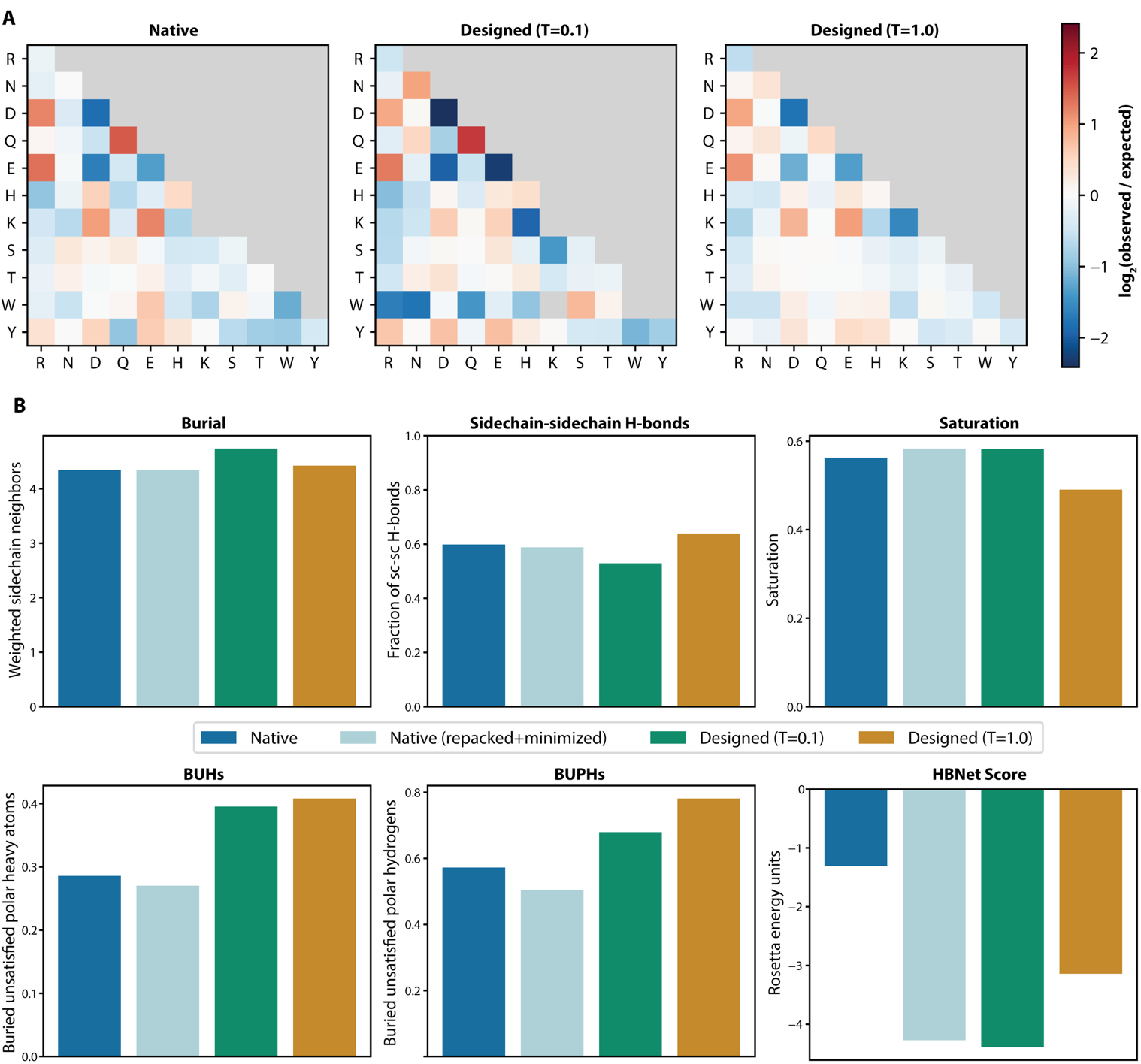

**Extended Data Figure 1: Native likeness of designed hydrogen bond networks. A)** Pairwise residue enrichment for native networks and networks generated by HBDesigner on the same randomly sampled backbones from the HBDesigner test set (n=10,000). Residue pairs are colored by log-enrichment (observed over expected co-occurrence from independent sampling). Redundant pairings or those with no observations are shaded grey. **B)** Selected structural metrics for native and designed networks sampled on the same backbones (mean value, n=1,000). Designs generated with low (T=0.1) and high (T=1.0) temperature are both included. Only successfully repacked designs forming one continuous network are included in the structural metrics.

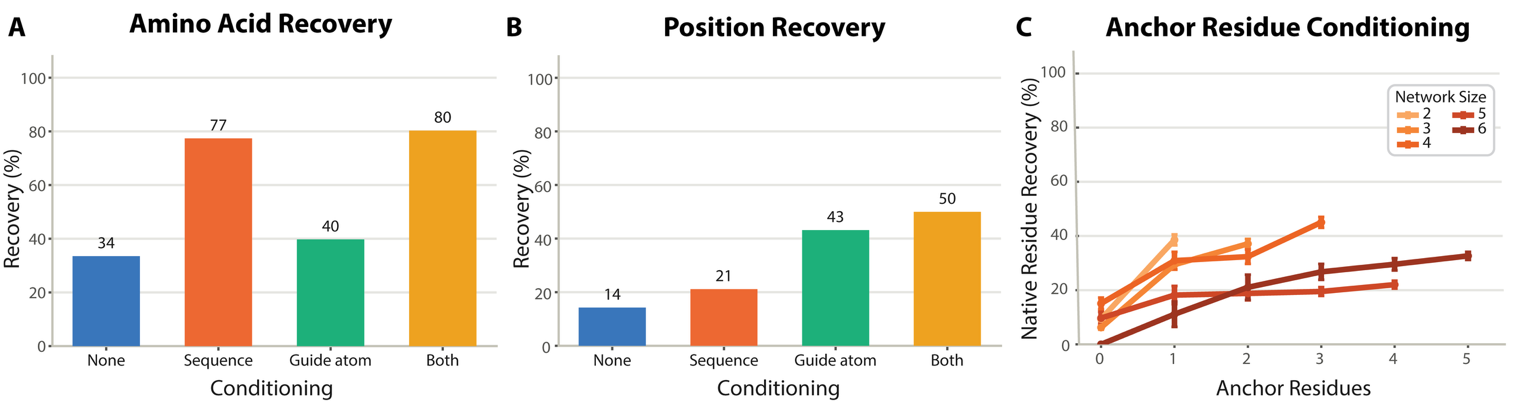

**Extended Data Figure 2: Model conditioning success rates.** Percent recovery of **A)** native network amino acids and **B)** native network positions, with different types of conditioning enabled. **C)** Percent recovery of native residues (correct position and amino acid) vs the number of anchor residues provided. Each line color represents a different total network size (including anchor residues). Each configuration is evaluated on randomly sampled native networks from the HBDesigner test set (n=1,000). For this experiment, sequence and guide atom conditioning are provided as soft constraints, while anchor residues are provided as hard constraints.

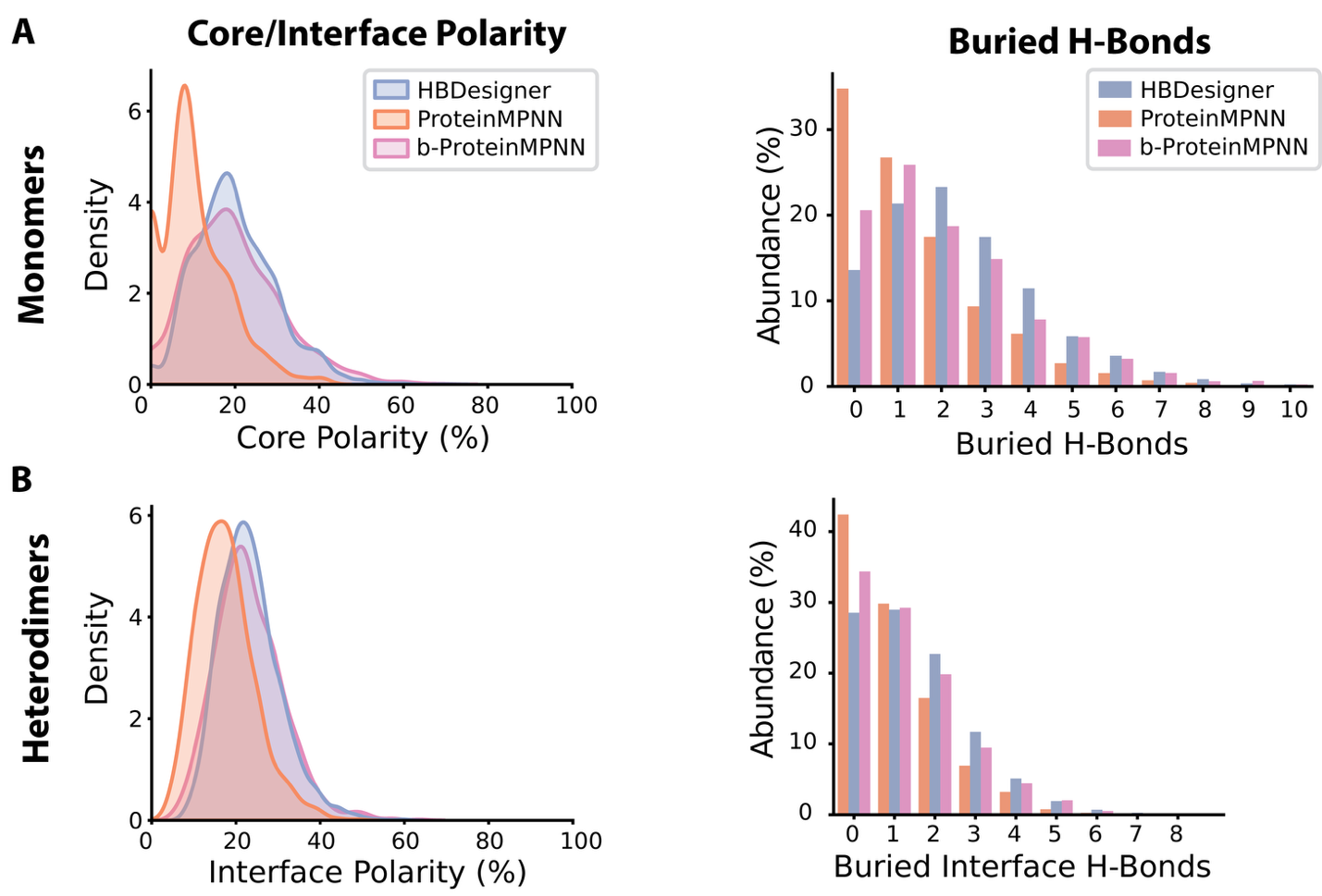

**Extended Data Figure 3: Additional refolding benchmark results.** **A)** Core polarity and buried core H-bonds for designed de novo monomers. **B)** Interface polarity and buried interface H-bonds for designed de novo heterodimers. Here, polarity is calculated as the percentage of polar residues present in a given region.

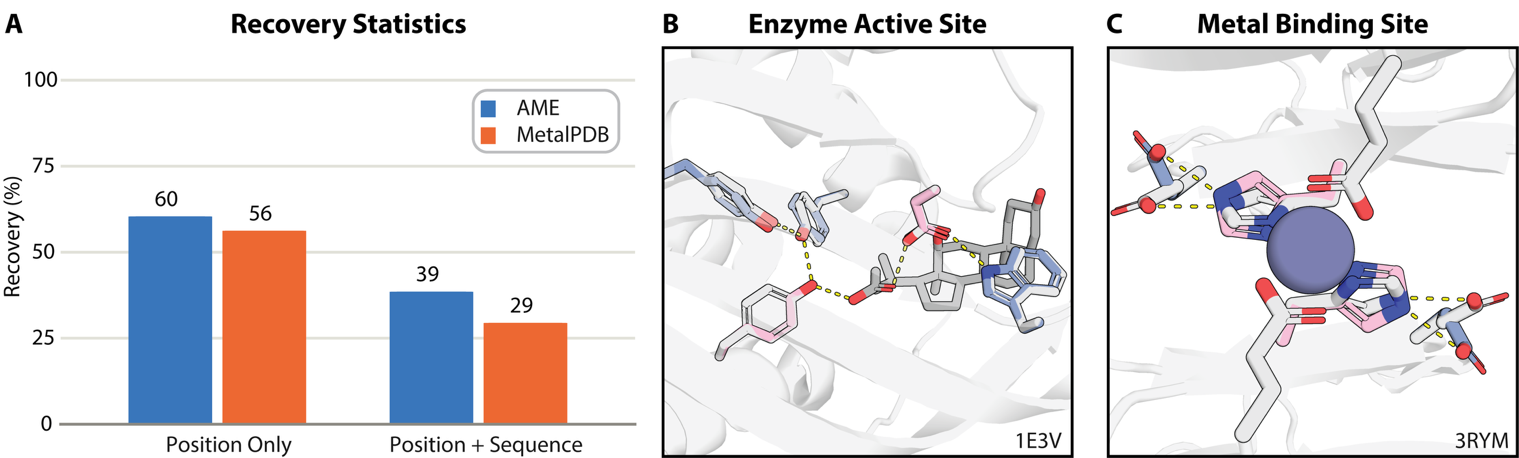

**Extended Data Figure 4: Second-shell network design benchmark. A)** HBDesigner native recovery statistics for second-shell interactions curated from the MetalPDB (67 sites, 112 designed residues) and AME (54 sites, 96 designed residues) datasets. Non-anchor residues present in the top-10 ranked designs for each site are scored as recovered. **B-C)** Example HBDesigner prediction successfully recovering second-shell contacts for a native **B)** enzyme active site and **C)** metal binding site.

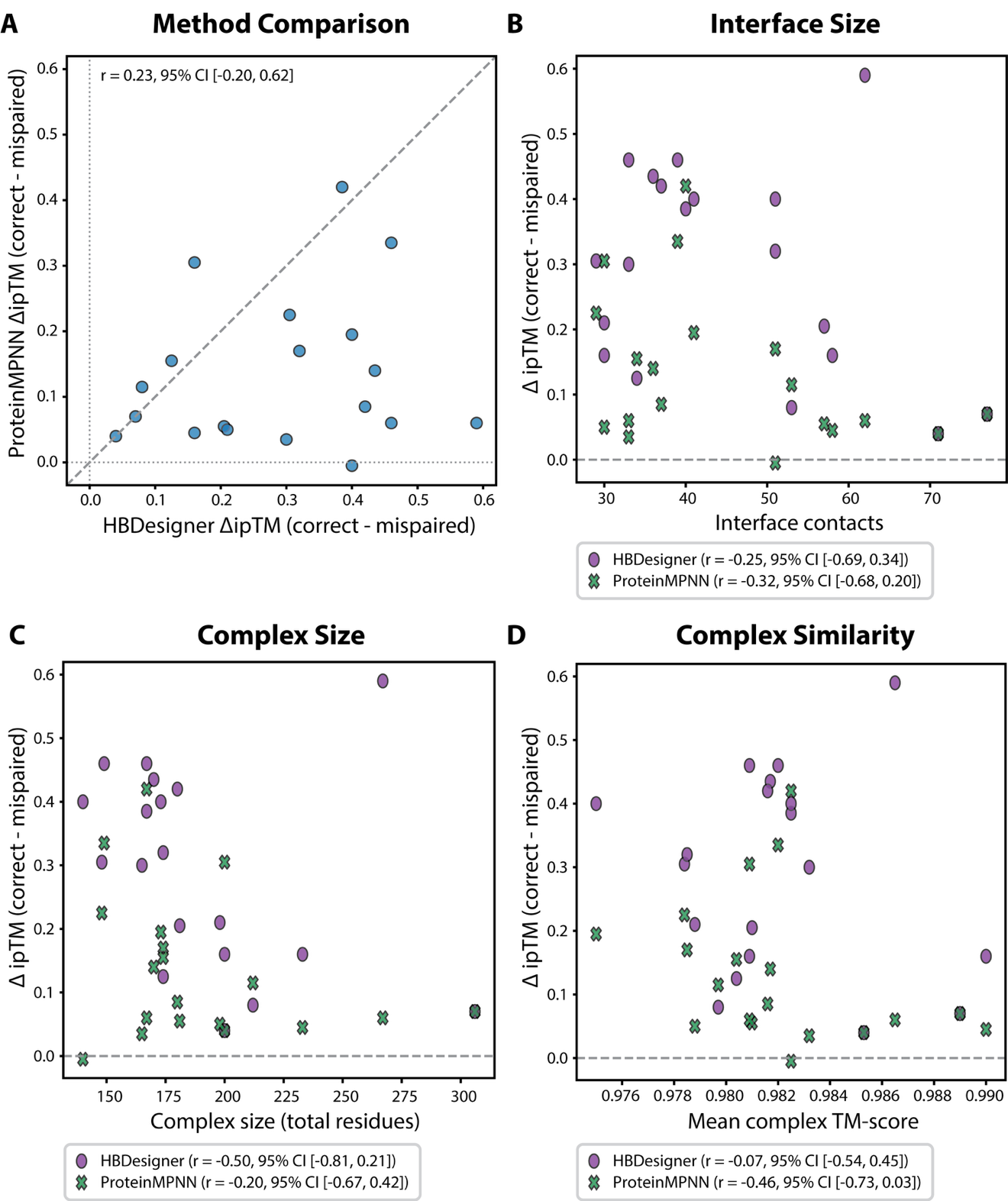

**Extended Data Figure 5: Family-level analysis of designed heterodimer specificity. A)** HBDesigner vs ProteinMPNN predicted design specificity (ΔipTM, AlphaFold3 ipTM of correctly-paired dimer minus median ipTM of mis-paired dimers). **B)** Interface size (Cα contacts within 8 Å), **C)** complex size, and **D)** complex similarity (TM-score) of each family vs ΔipTM. Interface size and complex were calculated on the geometric centroid of each family, and complex similarity was calculated for all family members against the centroid. Spearman correlation (r) is reported for each method with 95% confidence intervals obtained from 10,000 bootstrap samples.

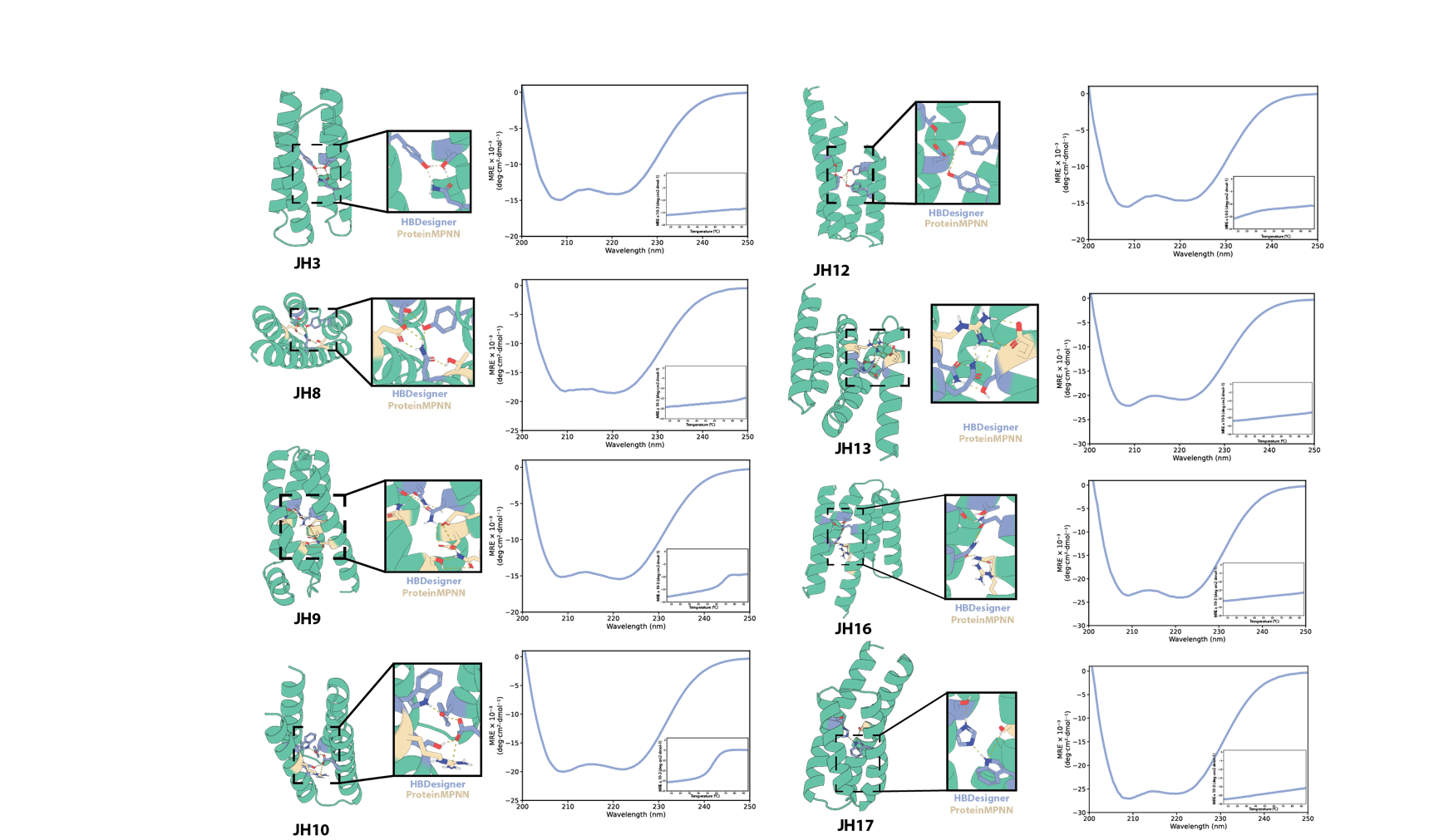

**Extended Data Figure 6: Characterization of additional monomers with highly polar cores.** Design models (left) of experimentally characterized monomers. Insets show designed H-bond networks comprised of residues introduced by HBDesigner (blue) and ProteinMPNN (tan). Circular dichroism (CD) spectra (right) of the selected monomers. Insets show CD temperature melts gathered over 0-95° C.

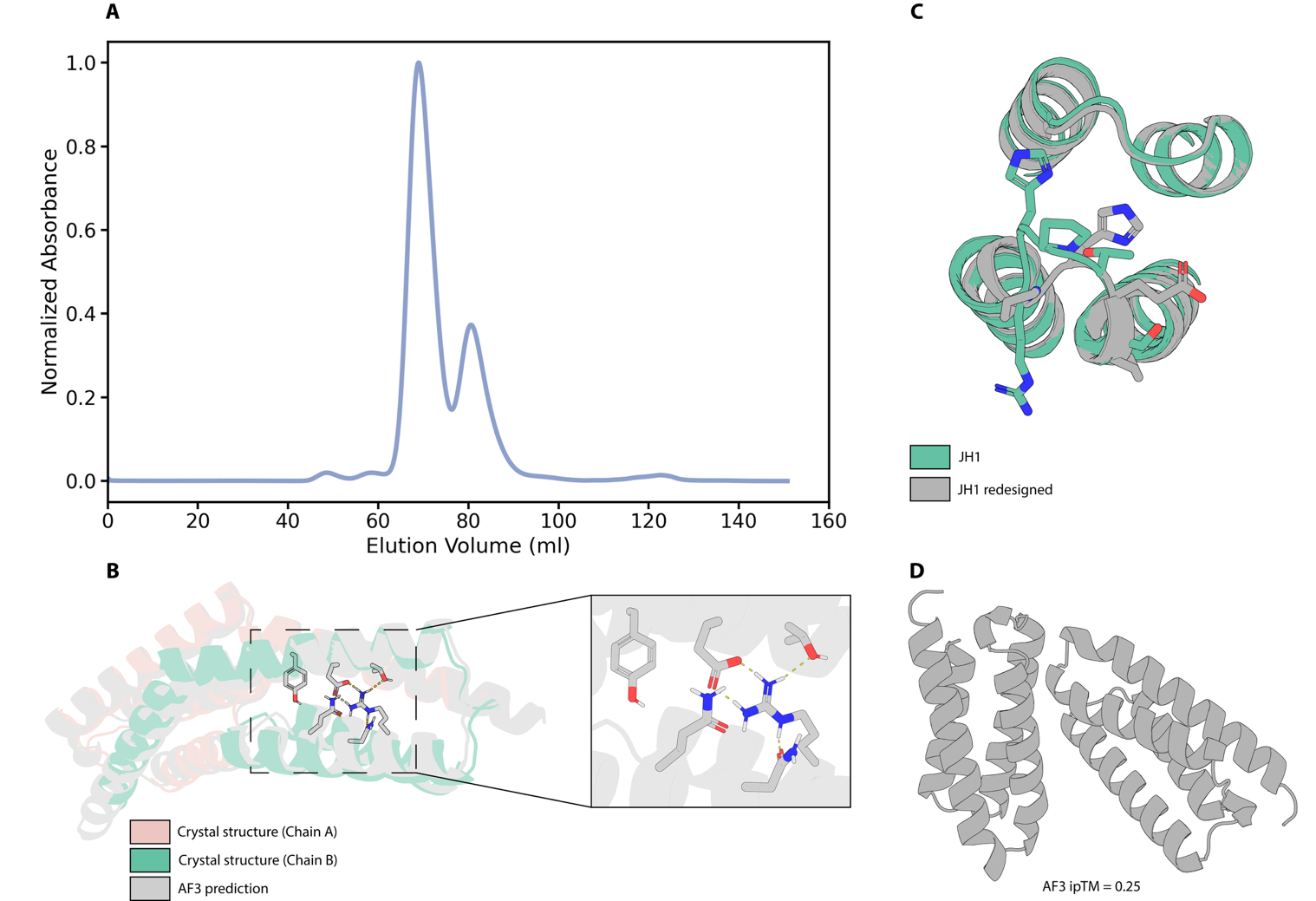

**Extended Data Figure 7: Additional characterization of design JH1. A)** Size exclusion chromatography (SEC) trace for design JH1. **B)** AlphaFold3 prediction of design JH1 as a dimer overlaid on the JH1 crystal structure (PDB: 13JM, 2.1 Å). **C)** Overlay of AlphaFold3 monomer predictions for design JH1, before and after loop redesign. **D)** AlphaFold3 dimer prediction for design JH1 after loop redesign.

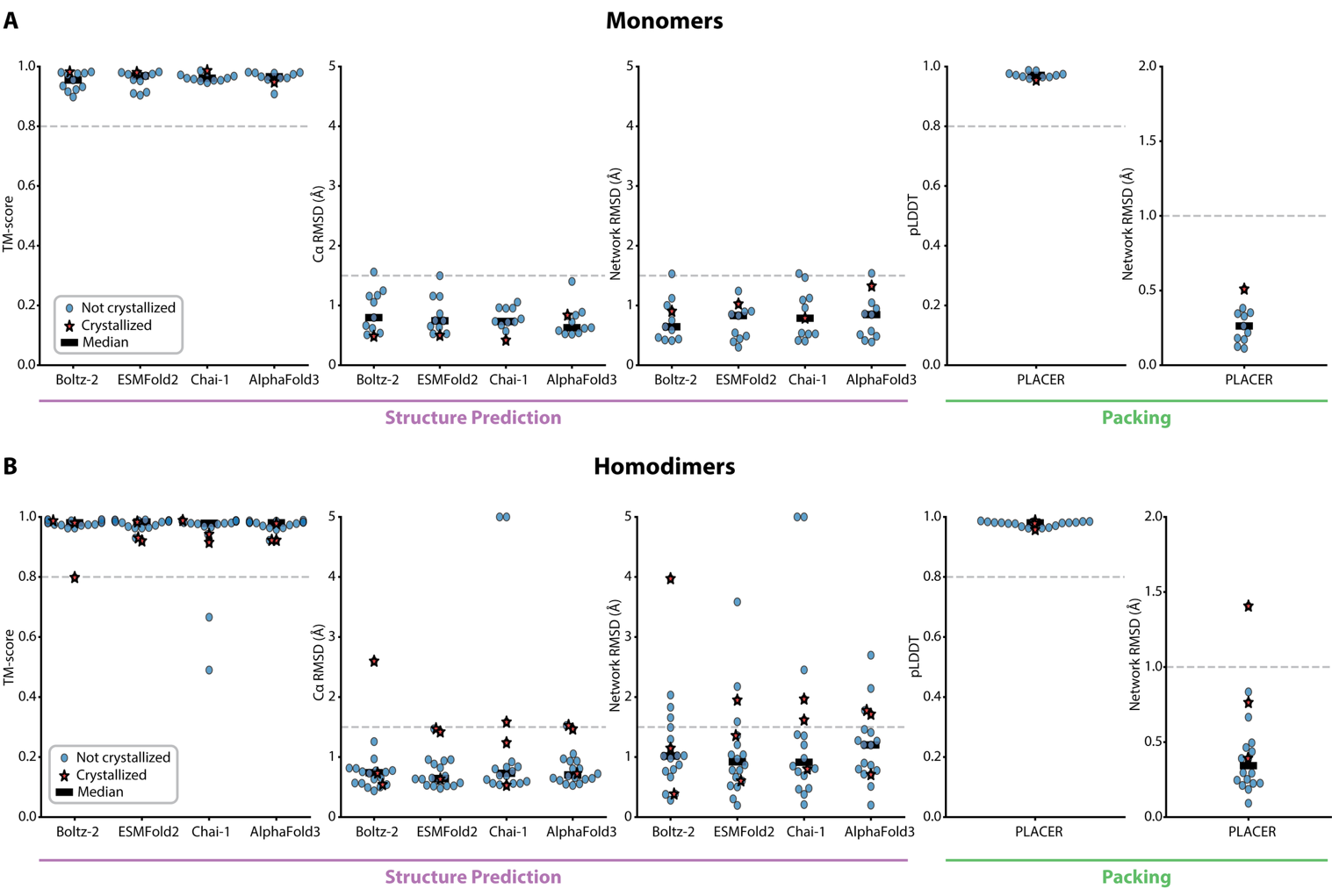

**Extended Data Figure 8: Computational structural validation.** Structure prediction (left) and packing (right) metrics for **A)** expressed and validated de novo monomer (n=11, including 1 crystallized) and **B)** de novo homodimers (n=18, including 3 crystallized). TM-score and RMSD metrics are calculated against the original HBDesigner design model. Note that RMSD values are clipped to a maximum of 5 Å for clarity.

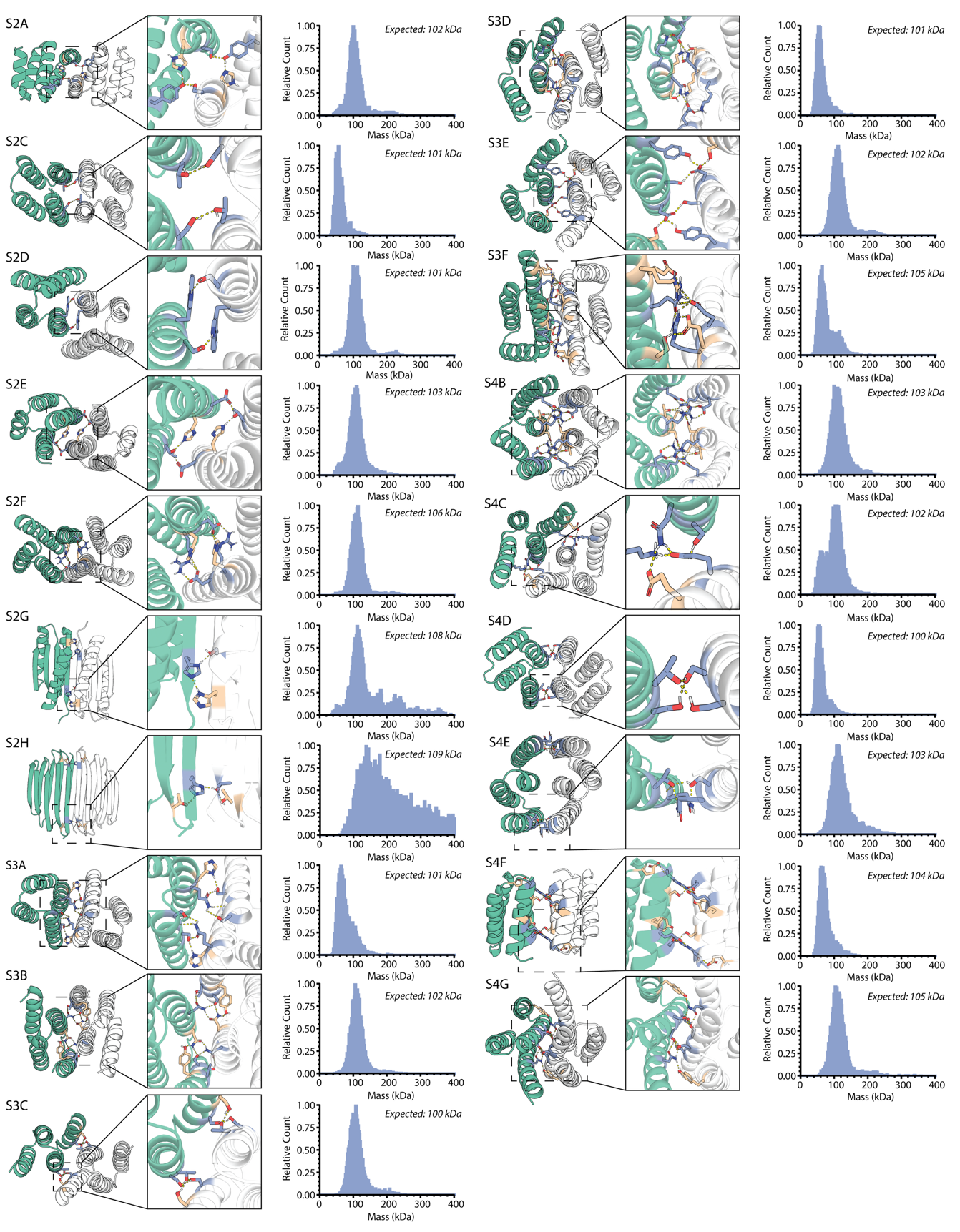

**Extended Data Figure 9: Characterization of additional homodimers with single interface hydrogen bond networks.** Design models (left) show homodimer chains in green and white. Details (center) show the designed networks with residues introduced by HBDesigner and LigandMPNN in blue and tan, respectively. Mass photometry plots (right) show oligomeric state for each homodimer labeled with maltose-binding protein.

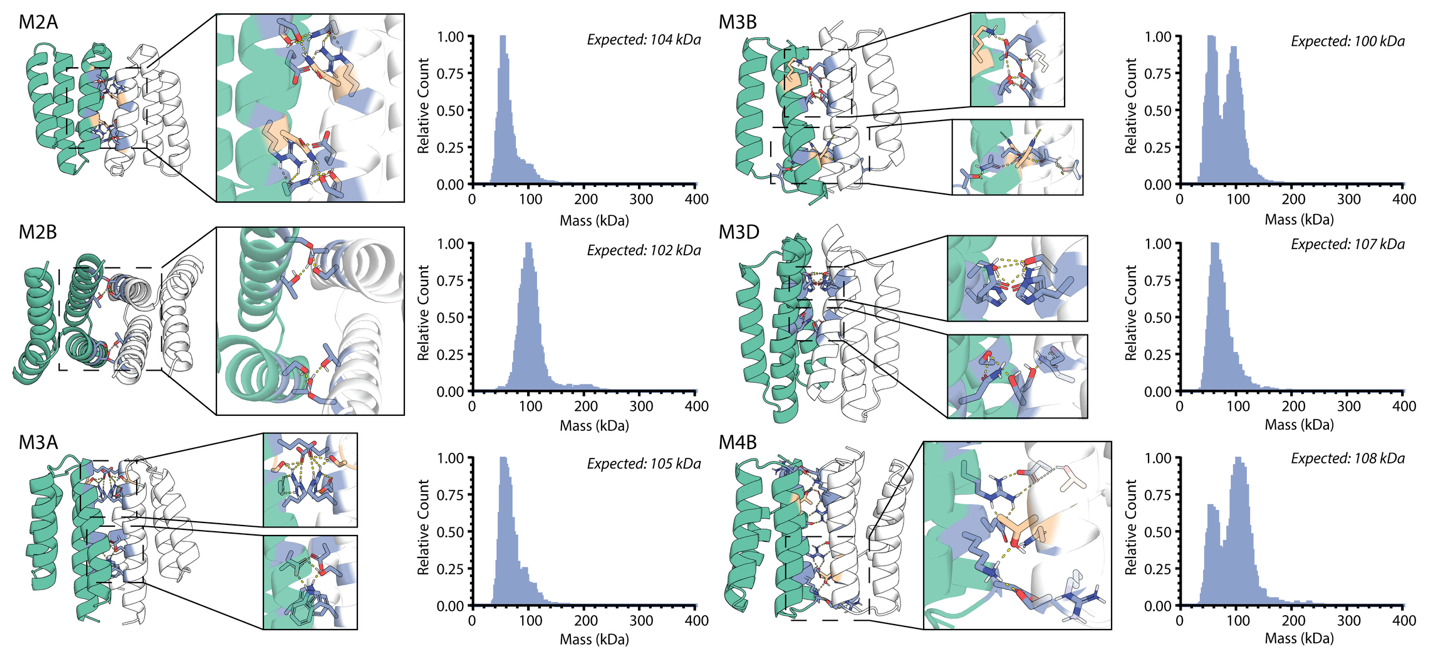

**Extended Data Figure 10: Characterization of additional homodimers with multiple interface hydrogen bond networks.** Design models (left) show homodimer chains in green and white. Details (center) show the designed networks with residues introduced by HBDesigner and LigandMPNN in blue and tan, respectively. Mass photometry plots (right) show oligomeric state for each homodimer labeled with maltose-binding protein.

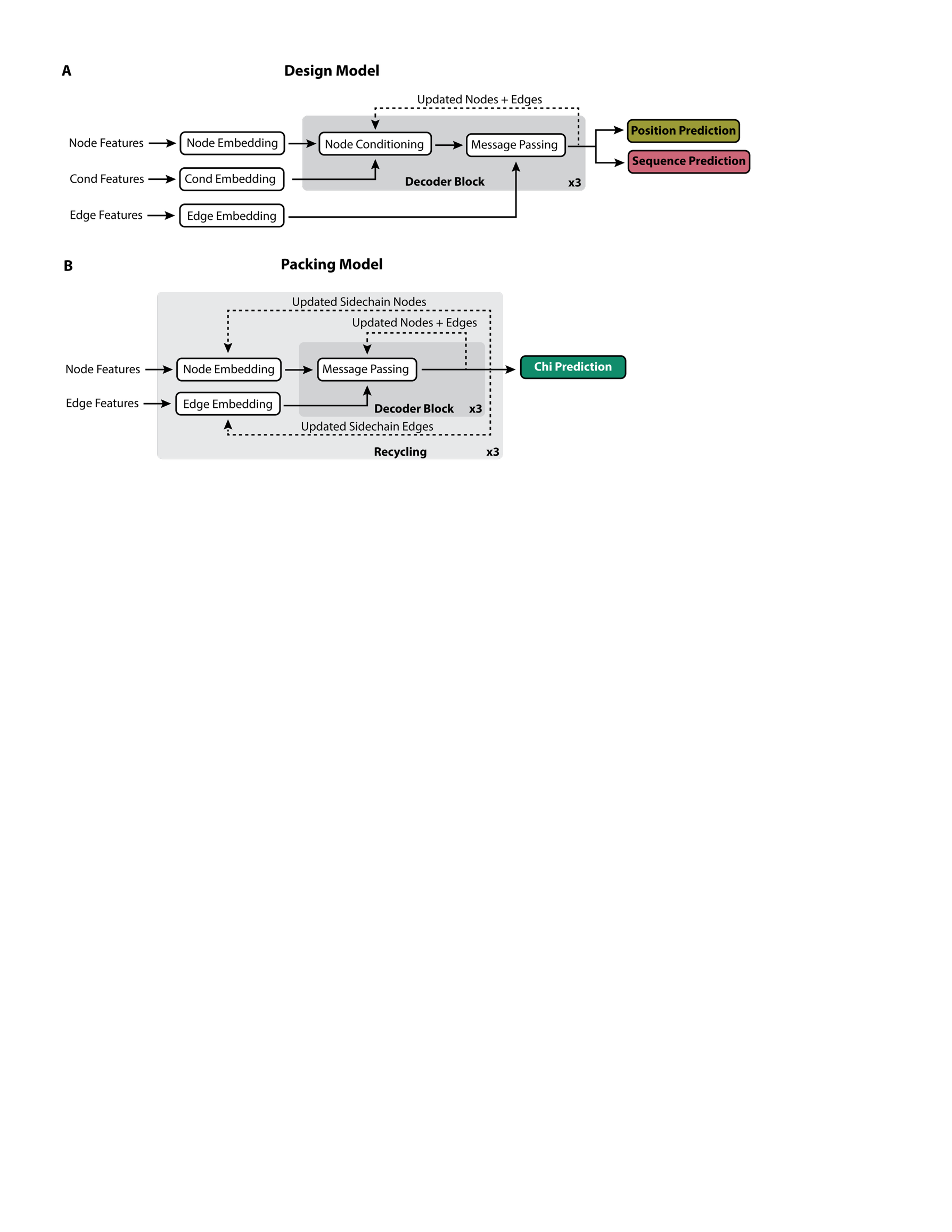

**Extended Data Figure 11: Architecture of the HBDesigner design and packing models.** **A)** The design model consists of three decoder blocks. In each block, nodes are updated with the conditioning information, then both nodes and edges are updated with message passing. Position and sequence predictions are calculated from the final node representations. **B)** The packing model uses three decoder blocks and three recycles. After each pass through the model, new sidechain angles and coordinates are recalculated and used to generate new nodes and edges. Gradients are only calculated on the final pass through the model.

**Supplementary Table 1: Description of input features for HBDesigner sequence design model.**

| **Type** | **Name** | **Shape** | **Description** |
| --- | --- | --- | --- |
| Node | bb_dihedral | [L, 6] | Sine and cosine encoding of backbone  ϕ, ψ, and ω dihedral angles. |
| Node | aatype | [L, 21] | One-hot encoding of amino acid types. All residues except previously decoded network residues are encoded as GLY. |
| Edge | atom_rbf | [E, 400] | Radial basis function encoding of pairwise distances between neighboring backbone atoms. |
| Edge | seq_sep | [E, 66] | One-hot encoding of relative sequence separation, or one bit to encode inter-chain neighbors. |
| Cond | net_res_num | [7] | One-hot encoding of number of network residues remaining to decode. |
| Node/Cond | guide_atom_rbf | [L, 16] | Radial basis function encoding of Cβ distance from each residue to the virtual guide atom. |
| Cond | guide_seq_dist | [21] | Probability distribution encoding the sequence conditioning. |

L=number of nodes, E=number of edges

**Supplementary Table 2: HBDesigner inference settings used for the refolding benchmark.**

| **Scaffold** | **Residues** | **Samples** | **Min Burial** | **Training Noise (Å)** |
| --- | --- | --- | --- | --- |
| Monomer | 2 | 200 | 4.0 | 0.2 |
| Monomer | 3 | 200 | 4.0 | 0.2 |
| Monomer | 4 | 500 | 4.0 | 0.02 |
| Monomer | 5 | 1000 | 4.0 | 0.02 |
| Monomer | 6 | 2000 | 4.0 | 0.02 |
| Heterodimer | 2 | 200 | 3.0 | 0.2 |
| Heterodimer | 3 | 200 | 3.0 | 0.2 |
| Heterodimer | 4 | 500 | 3.0 | 0.02 |
| Heterodimer | 5 | 1000 | 3.0 | 0.02 |
| Heterodimer | 6 | 2000 | 3.0 | 0.02 |

**Supplementary Table 3: Crystal data collection and refinement statistics.**

|  | JH1  (PDB ID: 13JM) | S2B  (PDB ID: 35SL) | S3G  (PDB ID: 13FC) | S4A  (PDB ID: 35SB) |
| --- | --- | --- | --- | --- |
| **Data Collection** |  |  |  |  |
| Resolution (Å) | 30.22 - 2.1 | 37.74 - 1.80 | 39.46 - 1.79 | 40.31 - 1.83 |
|  | (2.175 - 2.1) | (1.83 - 1.80) | (1.82 - 1.79) | (1.85 - 1.83) |
| Space group | P 21 21 21 | P 32 2 1 | P 21 21 21 | P 41 21 2 |
| a, b, c (Å) | 41.987 48.6667 87.0405 | 43.578 43.578 151.542 | 46.838 50.111 73.253 | 82.534 82.534 187.932 |
| α, β, γ(°) | 90 90 90 | 90 90 120 | 90 90 90 | 90 90 90 |
| R_Meas_ | 0.1232 (0.6636) | 0.055 (0.971) | 0.06714 (0.6691) | 0.1416 (2.769) |
| I / σ | 10.12 (3.35) | 21.5 (1.13) | 19.54 (4.19) | 9.68 (1.03) |
| Completeness (%) | 91.54 (93.14) | 99.0 (90.8) | 93.10 (93.90) | 99.90 (99.50) |
| Unique reflections | 9987 (977) | 29399 (1260) | 29241 (1377) | 109368 (4011) |
| Redundancy | 5.3 (5.2) | 9.0 (4.9) | 6.20 (5.96) | 13.17 (11.87) |
| CC1/2 | 0.998 (0.452) | 0.999 (0.633) | 0.998 (0.790) | 0.999 (0.438) |
| CC* | 0.999 (0.789) | 1 (0.831) | 0.999 (0.874) | 1 (0.628) |
| **Refinement** |  |  |  |  |
| Resolution (Å) | 30.22 - 2.1 | 37.74 - 1.80 | 39.46 - 1.79 | 40.31 - 1.83 |
| No. reflections | 9986 (978) | 16088 (680) | 16557 (730) | 58081 (2089) |
| R_Work_ / R_Free_ | 0.2273 / 0.2725 | 0.1843 / 0.2399 | 0.1682 / 0.2155 | 0.1956 / 0.2287 |
|  | (0.2896/ 0.3421) | (0.4251 / 0.4857) | (0.2872 / 0.3583) | (0.2284 / 0.3037) |
| No. atoms | 1551 | 1414 | 1511 | 5945 |
| Protein | 1530 | 1335 | 1438 | 5617 |
| Ligand/ion | 0 | 7 | 12 | 24 |
| Water | 21 | 72 | 71 | 304 |
| Ramachandran outliers (%) | 0.59 | 0 | 0 | 0 |
| Rotamer outliers (%) | 0 | 0 | 0 | 0 |
| Clashscore | 8.39 | 1.11 | 0.70 | 1.21 |
| B-factors (Å^2^) | 37.99 | 52.45 | 27.64 | 38.66 |
| Protein | 38.04 | 52.21 | 27.08 | 38.42 |
| Ligand/ion |  | 82.3 | 45.98 | 51.05 |
| Water | 34.47 | 54.11 | 35.93 | 42.11 |
| Bond lengths (Å) | 0.003 | 0.009 | 0.007 | 0.006 |
| Bond angles (°) | 0.54 | 0.93 | 0.8 | 0.78 |

* Highest resolution shell values shown in parenthesis.
